# Plasma membrane damage causes NLRP3 activation and pyroptosis during *Mycobacterium tuberculosis* infection

**DOI:** 10.1101/747014

**Authors:** Kai S. Beckwith, Marianne S. Beckwith, Sindre Ullmann, Ragnhild Sætra, Haelin Kim, Anne Marstad, Signe E. Åsberg, Trine A. Strand, Harald A. Stenmark, Trude H. Flo

## Abstract

*Mycobacterium tuberculosis* (Mtb) is a major global health problem and causes extensive cytotoxicity in patient cells and tissues. Here we define an NLRP3, caspase-1 and gasdermin D-mediated pathway to pyroptosis in human monocytes following exposure to Mtb. We demonstrate an ESX-1 mediated, contact-induced plasma membrane (PM) damage response that occurs during phagocytosis or from the cytosolic side of the PM after phagosomal rupture in Mtb infected cells. This PM injury in turn causes K+ efflux and activation of NLRP3 dependent IL-1β release and pyroptosis, facilitating the spread of Mtb to neighbouring cells. Further we reveal a dynamic interplay of pyroptosis with ESCRT-mediated PM repair. Collectively, these findings reveal a novel mechanism for pyroptosis and spread of infection acting through dual PM disturbances both during and after phagocytosis. We also highlight dual PM damage as a common mechanism utilized by other NLRP3 activators that have previously been shown to act through lysosomal damage.

**Graphical abstract:** 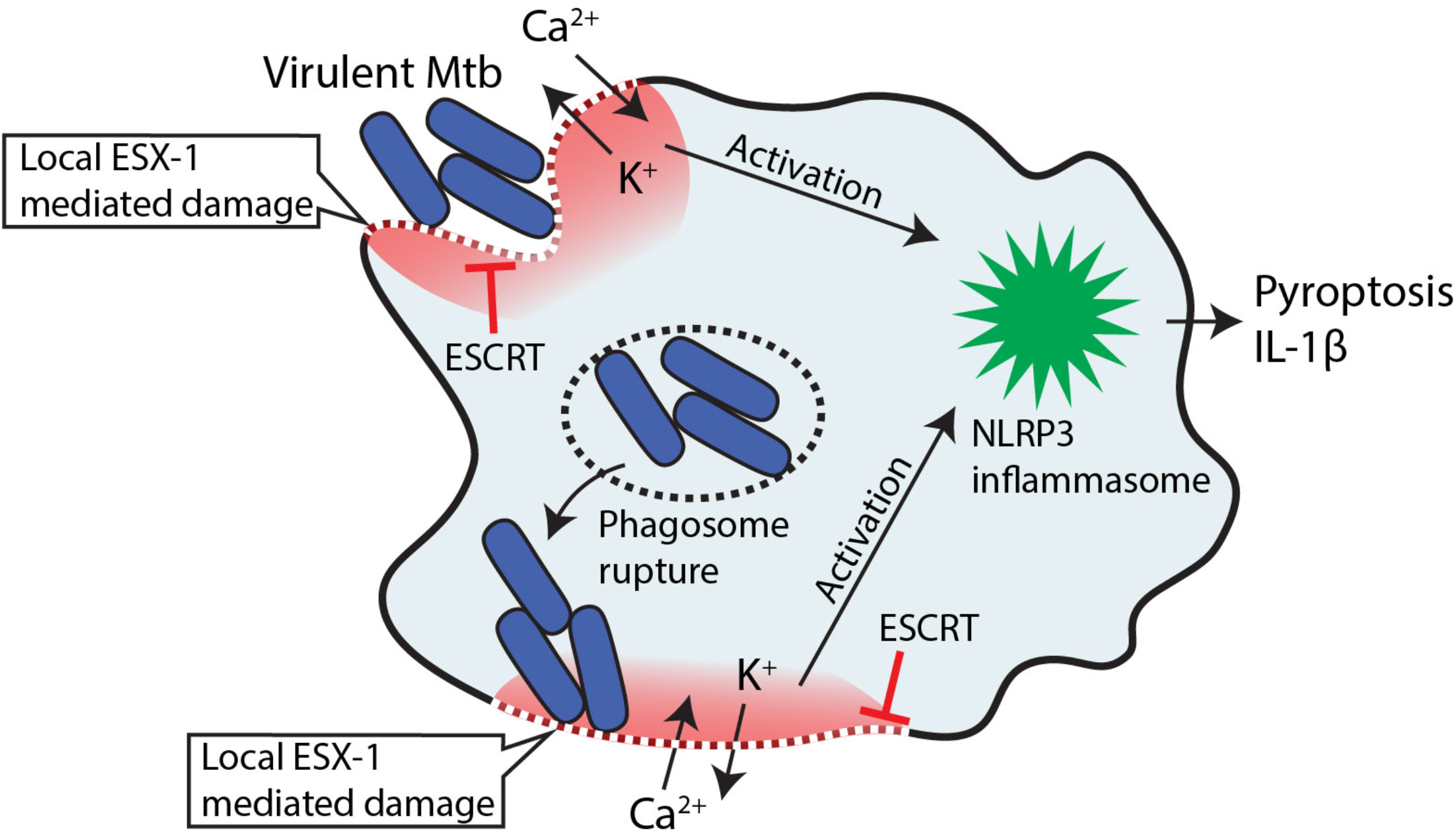

## Introduction

*Mycobacterium tuberculosis* (Mtb) is a deadly human pathogen, causing about 1,6 million deaths per year ^1^. A pathological hallmark of Mtb infection is extensive necrosis in infected tissues ^2^. Necrosis has long been regarded as an unregulated type of cell death, but recently several programmed necrotic pathways have been identified ^3, 4^. A highly inflammatory form of programmed necrosis is pyroptosis, occurring mainly in myeloid cells after pattern-recognition receptor (PRR) activation. In the classical pathway, activation of nucleotide-binding oligomerization domain-like receptors (NLRs) or absent in myeloma 2 (AIM2)-like receptors (ALRs) by pathogen- or self-ligands drives the macromolecular assembly of an inflammasome consisting of oligomerized NLRs or ALRs, the adaptor apoptosis-associated speck-like protein containing a CARD (ASC), and caspase-1 ^5–7^. Autocatalytic activation and cleavage of caspase-1 enables cleavage of pro-inflammatory cytokines interleukin (IL)-1β and IL-18, as well as the pore-forming molecule gasdermin D (GSDMD) ^8, 9^. IL-1β is released through GSDMD pores and in larger amounts during pyroptosis, the lytic cell death that often follows GSDMD pore formation ^10–13^.

Canonical NLRP3 and AIM2 inflammasome activation has been implicated in IL-1β release during Mtb infection in mouse and human macrophages ^14–19^. The agonist of AIM2 is double-stranded DNA ^20–22^ while the direct agonists of NLRP3 are not known. With few exceptions, two steps are required for NLRP3 activation: The priming signal involves increased expression of pro-IL-1β as well as inflammasome components such as NLRP3 itself, while the second signal is characterized by a range of cell damage events such as potassium (K+) and chloride (Cl-) efflux, mitochondrial dysfunction, metabolic changes, calcium fluxes, *trans*-Golgi disassembly and lysosomal damage ^7, 23, 24–31, 32^. K+ efflux in particular is considered a key determinant of NLRP3 activation for a range of triggers ^28^ with some exceptions involving mitochondrial disruption and mtROS production, e.g. by the small-molecule compound Imiquimod ^29^. Mitochondrial dysfunction seems closely tied to Mtb-induced necrosis as well ^33, 34^, but whether this is related to inflammasome activation is not clear. Inflammasome activation and pyroptosis by Mtb is dependent on the type VII secretion system 6-kDa early secretory antigenic target (ESAT-6) secretion system 1, ESX-1. ESX-1 secretes a range of protein substrates, and is postulated to mediate phagosome permeabilization with full or partial translocation of Mtb into the cytosol ^35–38^. Phagosomal permeabilization presumably allows the release of Mtb DNA and activation of AIM2 ^19^, and release of active cathepsin B from damaged phagolysosomes has been proposed as a trigger for NLRP3 ^24, 39, 40^. However, off target inhibitor effects and negative results in knockout experiments have left the jury open on the role of cathepsin release in NLRP3 inflammasome activation ^28, 41–43^.

Here, we use time-lapse- and correlative microscopy to spatiotemporally resolve Mtb-induced inflammasome activation and cell death at a single-cell level. We show that contact-mediated PM damage caused by ESX-1 activity occurring either during phagocytosis, or from the cytosol after phagosomal escape, drives K+ efflux and subsequent NLRP3 activation and GSDMD-dependent pyroptosis of human macrophages. This mechanism is not unique to Mtb: Our findings identify a common mechanism where PM damage caused by Mtb or crystals triggers inflammasome activation, IL-1β release and pyroptosis unless the membrane damage is balanced by repair mediated by the endosomal sorting complexes required for transport (ESCRT) machinery ^44^.

## Results

### Mtb H37Rv infection induces canonical NLRP3 activation followed by pyroptosis

Based on previously published findings ^15–19^, we hypothesized that Mtb induces the assembly of one of the ASC-dependent inflammasomes, NLRP3 or AIM2, upon infection of macrophages. THP-1 macrophages expressing ASC-GFP were infected with Mtb H37Rv constitutively expressing BFP (Mtb::BFP) and imaged after 24 hours or by time-lapse microscopy. The cell-impermeable DNA dye DRAQ7 was present in the medium to assess cell death. ASC-GFP is evenly distributed in the cytosol and appears as diffuse green until inflammasome activation induces ASC aggregation, visible as bright specks ^45^. Mtb induced ASC speck formation in infected macrophages (Figure 1a and b) and specks were exclusively localized to dead cells as indicated by DRAQ7-positive nuclei. The number of dead cells, ASC specks and IL-1β secretion increased with increasing multiplicity of infection (MOI, Figure S1a). Treatment with the specific NLRP3 inhibitor MCC950 ^46^ or increased extracellular concentrations of KCl inhibited the formation of ASC specks, demonstrating that Mtb infection results in potassium efflux-driven activation of the NLRP3 inflammasome (Figure 1b). Correspondingly, IL-1β secretion was inhibited by MCC950 and KCl as well as the caspase inhibitors Z-VAD-FMK (pan-caspase) or VX765 (caspase-1). Treatment with Z-VAD, VX765, MCC950 or KCl all reduced cell death measured after 24 hours. Knocking down NLRP3 using CRISPR-Cas9 similarly reduced Mtb-induced cell death and completely inhibited ASC speck formation (Figure 1c and Figure S1b). Similar inhibitor- and NLRP3 KD results were obtained with the canonical NLRP3 inflammasome-inducing combination LPS and nigericin (Figure S1c-d). NLRP3-dependent IL-1β release and cell death was verified in Mtb-infected primary human monocytes (Figure 1d), indicating the physiological relevance of these findings. Live-cell time-lapse microscopy of single Mtb-infected cells revealed a nuclear DRAQ7 signal within few minutes after ASC speck formation, while the DRAQ7 signal remained low after ASC speck formation in THP-1 cells deficient in GSDMD (Figure 1e, Figure S1b). Together our data demonstrate that Mtb induces potassium efflux-driven activation of the canonical NLRP3 inflammasome upon infection of human monocytes and macrophages, followed by GSDMD-dependent pyroptotic cell death with release of IL-1β. The results also confirm ASC speck formation as a representative readout for inflammasome activation by Mtb.

**Figure 1.**
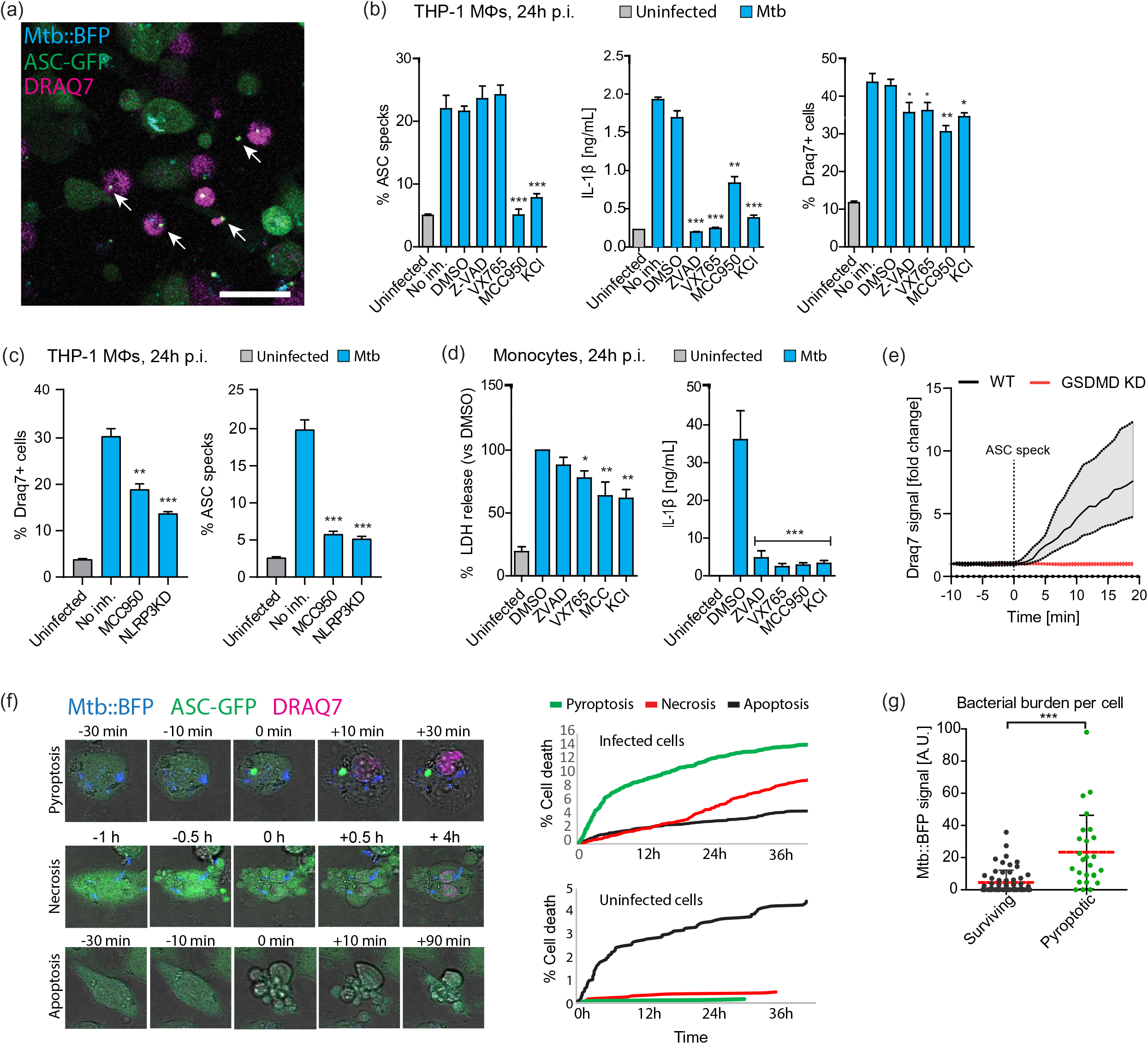
Mtb H37Rv infection induces canonical NLRP3 activation followed by pyroptosis. THP1 ASC-GFP macrophages were infected by MtbH37Rv::BFP at MOI 20, and imaged by time-lapse microscopy or at 24 hours post infection (p.i.). (**a**) ASC-GFP specks observed as bright green specks (white arrows), while diffuse cytosolic green indicates live ASC-GFP expressing THP1 cells. DRAQ7 binds DNA in the nuclei of dead cells (magenta). (**b**) THP1 ASC-GFP cells were treated with DMSO, Z-VAD-FMK (50µM), VX765 (50µM), MCC950 (10µM) or KCl (40mM) and infected with Mtb at MOI 20. DR+ cells and ASC specks were quantified at 24h p.i. (n>2000 cells in triplicates per condition). IL-1β release was determined by ELISA. Bars show mean±SEM. (**c**) THP1 ASC-mNeonGreen wild-type or NLRP3 knockdown cells were infected by Mtb and imaged 24h p.i. (**d**) Primary human monocytes were treated with inhibitors and infected with Mtb as before, and LDH and IL-1β release were determined 24h p.i. Bars show mean±SD for 4 donors. (**e**) THP1 ASC-mNeonGreen wild-type or GSDMD knockdown cells were infected with Mtb and imaged by time-lapse microscopy. DRAQ7 intensity during ASC speck formation was measured in single cells in WT (n=57) and GSDMD KD (n=81) cells. Median±IQR shown. (**f**) The cumulative number of pyroptotic, necrotic and apoptotic cell death events in THP1 ASC-GFP cells was quantified from a 40h time-lapse experiment during infection with Mtb. Representative cell death events in each class are shown. (**g**) The bacterial burden (intracellular Mtb::BFP fluorescence) was determined immediately prior to ASC speck formation (n=26), and at the average time of ASC speck formation, in cells surviving a 24h time-lapse experiment (n=61). Lines indicates median values. *p<0.05, **p<0.01, ***p<0.001, ****p<0.0001, by one-way ANOVA with Dunnett’s test in (a)-(d), and Mann-Whitney test in (g). Data in (a)-(e) representative of 3-5 independent experiments. See also Figure S1.

To gain a better understanding of the kinetics of pyroptosis caused by Mtb, we imaged THP1-ASC-GFP cells infected with Mtb-BFP for 40 hours by time-lapse microscopy. We noted the cumulative number of cell deaths by pyroptosis, apoptosis and necrosis, scored as illustrated in Figure 1f and corresponding Movies S1, S2 and S3. Pyroptotic cells were characterized as those forming a visible ASC speck followed by rapid influx of DRAQ7, while necrotic cells did not form ASC specks and generally followed a slower progression with e.g. cessation of cell migration prior to a slower influx of DRAQ7. Apoptotic cells were identified by a characteristic blebbing morphology and no DRAQ7 influx. Pyroptosis was the dominating form of cell death during the first 24 hours of infection. In line with this result, the necroptosis inhibitors Nec1s (RIPK1 inhibitor) and GSK’872 (RIPK3 inhibitor) did not affect cell death or ASC speck formation after 24 hours of infection (Figure S1e). Pyroptosis and necrosis primarily occurred in infected cells, while the proportion of apoptotic cell death was larger in uninfected bystander cells (Figure 1f). Additionally, the bacterial burden was higher in cells that formed ASC specks than in infected cells that did not (Figure 1g). These results indicate that pyroptosis from Mtb infection is a cell-intrinsic effect, i.e. a direct result of infection within single cells.

We reasoned the relative reduction in pyroptosis and increase in necrosis from 24-48 could be due to prolonged stimulation of Toll-like receptor (TLR) signalling, which has been reported to inhibit NLRP3 inflammasome activation ^47^. Indeed, we observed that prolonged treatment (24h) of THP-1 cells with TLR2 or 4 ligands prior to infection with Mtb decreased cell death and ASC speck formation to a similar extent as treatment with MCC950 (Figure S1f).

### Pyroptosis causes severe cellular damage and allows for spread of Mtb

Cell death is accompanied with morphological changes that can inform about which processes are involved in execution of the particular cell death pathway. We investigated the ultrastructural features of pyroptosis by correlative 3D light and electron microscopy (CLEM). Focused Ion Beam -Scanning EM (FIB-SEM) was used to gain ultrastructural information with near-isotropic 3D resolution, using a correlation system we developed previously ^48^. Uninfected THP1-ASC-GFP cells displayed a normal, healthy morphology (Figure 2a). In comparison, infected, pyroptotic cells (i.e. with visible ASC specks) had severe ultrastructural disruptions, including fragmented and occasionally split nuclear membranes, large PM disruptions and leakage of content to the extracellular space. Most organelles were no longer recognizable, with the exception of mitochondria that could be discerned by their internal cisterna. The mitochondria displayed a more rounded and less interconnected morphology than in the healthy control cell. Mtb displayed a largely intact morphology, and some bacteria were observed in contact with the extracellular space, suggesting a means of escape from pyroptotic cells. Cells that were treated with LPS and nigericin and formed ASC specks displayed similar ultrastructural features as Mtb-infected pyroptotic cells, with nuclear and PM disruptions, cytosol leakage and organelle damage (Figure S2a).

**Figure 2.**
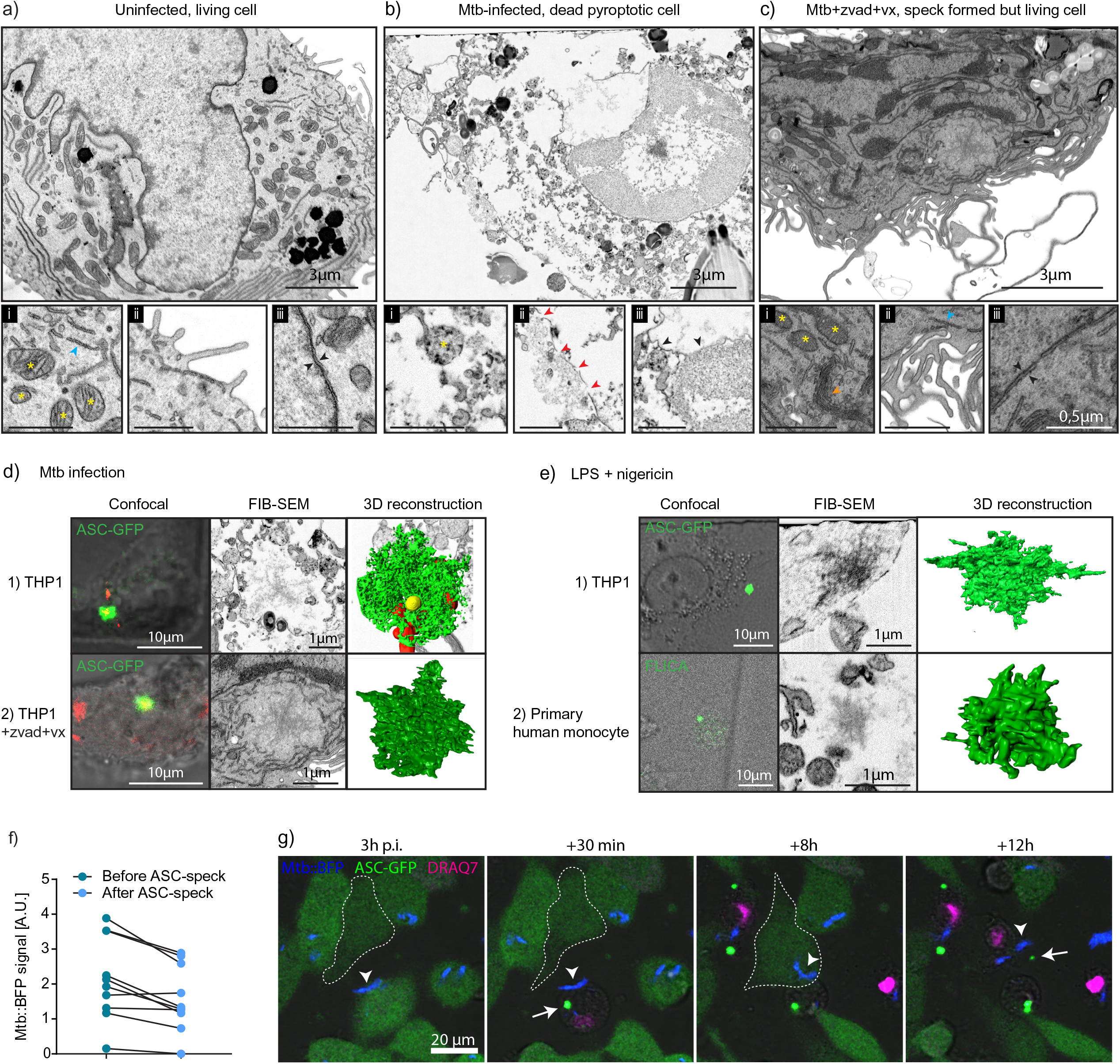
Pyroptosis causes severe cellular damage and allows for spread of Mtb. Ultrastructural comparison by FIB-SEM tomography between (**a**) uninfected (**b**) infected, pyroptotic and (**c**) infected, living cell with an activated inflammasome. THP1 ASC-GFP cells were infected with Mtb H37Rv::BFP (MOI 20) and fixed after (**b**) 24 hours or (**c**) 5h in the absence or presence of Z-VAD FMK (50µM) and VX765 (50µM), respectively. The uninfected cell (**a**) is from a 24h infection experiment but was found to be uninfected from confocal microscopy and FIB-SEM. Insets (i), (ii) and (iii) highlight mitochondria/ER/Golgi, plasma membrane morphology and nuclear membrane morphology, respectively. Yellow stars indicate mitochondria, blue arrowheads ER, orange arrowheads Golgi, red arrowheads indicate plasma membrane ruptures and black arrowheads indicate nuclear membranes. Black inset scalebars 1µm. (**d**) Structure of ASC specks induced by Mtb infection in THP1 ASC-GFP cells. Cell number 1) and 2) are the same as in b) and c), respectively. Cells were imaged by correlative confocal microscopy and FIB-SEM, and ASC specks were reconstructed in 3D from the respective FIB-SEM image stacks. (**e**) Confocal and FIB-SEM correlative microscopy of ASC specks in THP1 ASC-GFP cells and primary human monocytes primed for 3h by LPS (10ng/ml) and treated with 6.7µM nigericin for 1-2h. Monocytes were additionally treated with FLICA caspase-1 reagent (FAM-YVAD-FMK, 20µM) for visualization of active caspase-1 at the inflammasome. (**f**) Quantification of intracellular Mtb::BFP fluorescence in single cells before and after ASC speck formation and pyroptosis for n=10 representative cells. (**g**) THP1 ASC-GFP (green) cells in DRAQ7 containing medium (magenta) and infected with Mtb::BFP (MOI 20, blue) imaged by time-lapse confocal microscopy for 24h. Arrowheads indicate Mtb that is phagocytosed twice during the time course, and arrows indicate ASC specks. See also Figure S2.

To distinguish morphological changes that occurred as a consequence of pyroptosis from those caused by Mtb infection prior to or during inflammasome activation, we infected cells for 5h with Mtb-BFP in the presence of the caspase inhibitors Z-VAD and VX765. Cells were imaged live during infection with DRAQ7 in the medium, and chemically fixed within two minutes of the last captured frame. Cells with an ASC speck but without influx of DRAQ7 and no visible changes to cell morphology were chosen for further imaging by FIB-SEM. The morphology was similar to uninfected cells, with intact organelles (Golgi, ER, interconnected and elongated mitochondria), a continuous PM and a double nuclear membrane (Figure 2c). Similar results were observed in THP-1 GSDMD KD cells (Figure S2b). Hence, the dramatic morphological changes in pyroptotic cells are a consequence of the pyroptotic cell death process itself.

Next, we investigated the structure of the ASC speck using the correlative imaging approach. ASC Specks were visible in SEM images as aggregates with a branched structure and a size of about 1 - 2,5 µm in THP1-ASC-GFP cells, both when infected with Mtb and when treated with LPS and nigericin (Figure 2d-e). The structure corresponds well to that previously observed in a zebrafish model ^49^. The speck in the living (Z-VAD + VX765-treated) cell was located in an open space in the cytosol of the cell, seemingly devoid of other organelles. As a control that the observed ASC speck structure was not a consequence of ASC overexpression or tagging, we treated primary human monocytes with LPS and nigericin, and visualised active caspase-1 by FAM-YVAD-FMK (FLICA). Caspase-1 is stabilized in its active form on the inflammasome ^50^, and we therefore hypothesized that the point with highest fluorescence intensity from FLICA treatment would be the point where the speck was assembled. Indeed, by CLEM, we could identify a similar branched structure in LPS and nigericin-treated primary human monocytes, albeit with a smaller size than that observed in THP-1 cells (about 0,8 µm). To our knowledge, this is the first time an ASC speck has been visualized in a primary human cell, without tagging or overexpression of selected proteins.

To elaborate on the possible consequences of pyroptosis on Mtb spreading as indicated by EM, we investigated pyroptosis during time-lapse imaging. On average, about 25% of Mtb bacilli were lost into the medium immediately after pyroptosis (Figure 2f), while the rest were trapped in the pyroptotic cell. However, Mtb that were trapped in pyroptotic cells could be phagocytosed by neighbouring cells. Further, this could induce ASC speck formation with pyroptosis in the new host cell, thus propagating inflammasome activation and cell death (Figure 2g, Movie S4). This indicates that pyroptosis allows some bacteria to spread immediately after cell death, and that bacteria that are trapped in the ghost cell can also contribute to further local spread of infection by efferocytosis.

### Mtb-induced inflammasome activation is dependent on ESX-1 but not on the tuberculosis-necrotizing toxin (TNT) or destabilization of host mitochondria

It has been reported that Mtb deleted in the Region of Difference 1 (MtbΔRD1), where most ESX-1 related genes are located, are deficient in inflammasome activation ^15–17^. Our results are in line with these reports, as MtbΔRD1 caused significantly less ASC speck formation and cell death of THP-1 cells than Mtb wild-type (Figure 3a). We also observed that Mtb grown in the absence of the detergent tween-80 are more potent in inflammasome activation than Mtb grown in the presence of tween (our regular growth condition). Detergent has been published to perturb the mycobacterial capsule, in particular decreasing the abundance of ESX-1 related proteins on the Mtb surface/capsule ^51, 52^. Our data thus show that inflammasome activation correlates with ESX-1 activity and the integrity of the Mtb capsule, suggesting that deposition of ESX-1 related factors on the Mtb surface/capsule is important for Mtb inflammasome activation.

**Figure 3.**
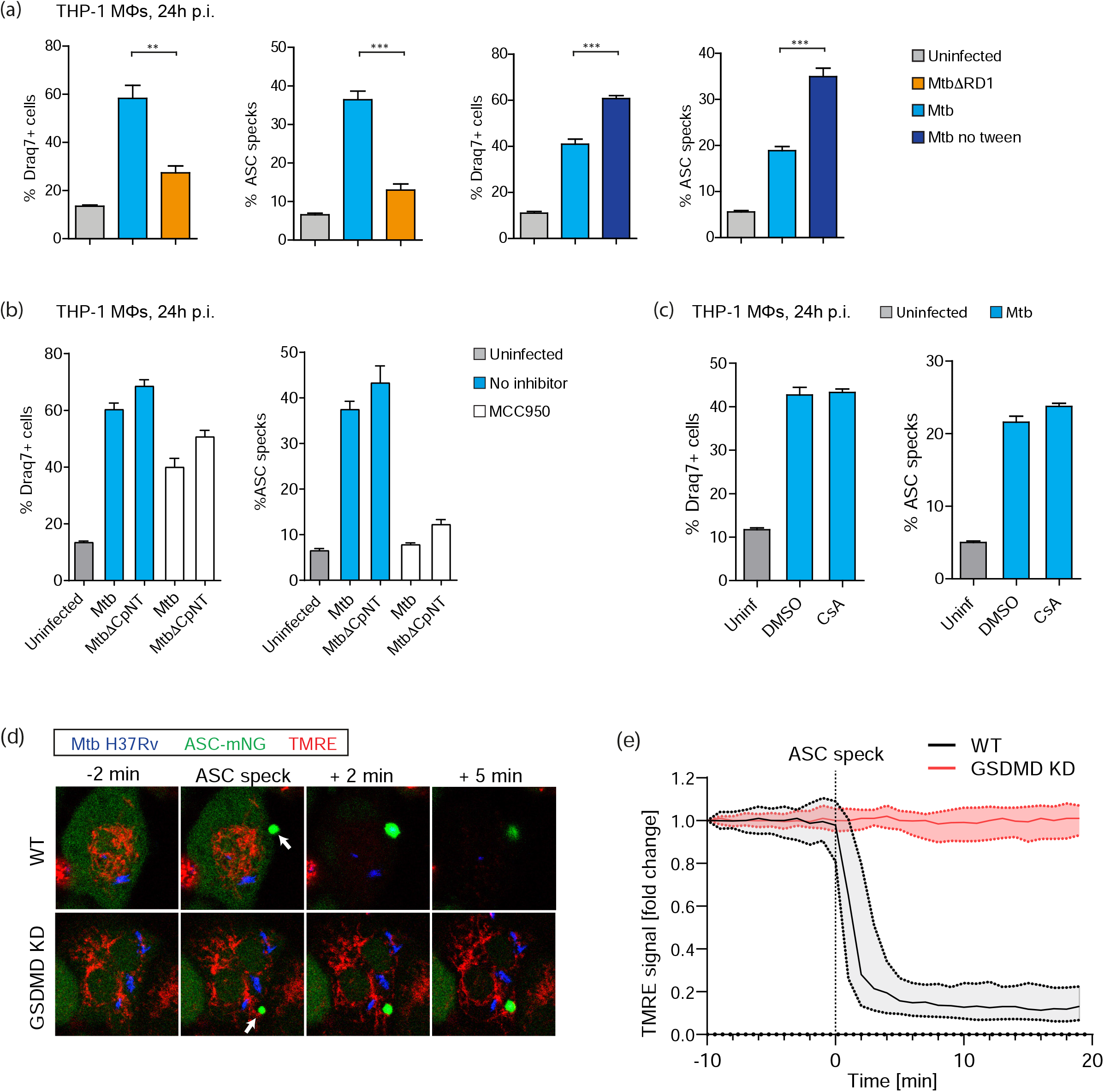
Mtb-induced inflammasome activation is dependent on ESX-1 but not on TNT or destabilization of host mitochondria. (**a**) THP1 ASC-GFP macrophages were infected with Mtb H37Rv ΔRD-1 or Mtb H37Rv cultured in the presence or absence of tween-80 at MOI 20. (**b**) THP1 ASC-GFP cells were not treated or treated with MCC950 (10µM) and infected with Mtb H37Rv or Mtb H37RvΔCpNT (MOI 20). (**c**) THP1 ASC-GFP cells were treated with DMSO or cyclosporine A (CsA, 5µM) and infected by Mtb H37Rv MOI 20. In (**a-c**) ASC specks and DR+ cells were quantified for n>2000 cells per condition, *p<0.05, **p<0.01, ***p<0.001, by one-way ANOVA with Dunnett’s test. Bars show mean±SEM. Data representative of 3 independent experiments. (**d**) Representative time-lapse images of THP1 ASC-mNeonGreen (green) cells or THP1 ASC-mNeonGreen cells depleted of GSDMD by CRISPR-Cas9, with mitochondria stained by TMRE (red, 10nM) and infected with Mtb::BFP (blue) during ASC speck formation. (**e**) Quantification of TMRE intensity, indicative of mitochondrial membrane potential before and after ASC speck formation in WT and GSDMD depleted cells (n=57 and n=81 cells). Median±IQR shown. Representative of 2 independent experiments. See also Figure S3.

Tuberculosis necrotizing toxin (TNT, the C-terminal end of the protein CpnT) is secreted from Mtb and released into the host cell cytosol in an ESX-1 dependent manner, leading to macrophage necroptosis ^34, 53, 54^. We therefore wanted to investigate if TNT is contributing to the ESX-1 dependent inflammasome activation. To this end, we infected THP1-ASC-GFP macrophages with wildtype Mtb, MtbΔCpNT or MtbΔCpNT complemented either with the catalytically active or inactive form of TNT (MtbΔCpNT::TNT and MtbΔCpNT::TNT* respectively). We found that all Mtb mutants and complemented strains were similarly competent in inflammasome activation, and that treatment with MCC950 had a similar effect across bacterial strains in inhibiting ASC speck formation (Figure 3b and Figure S3a). This indicates that TNT does not play a role in Mtb-induced inflammasome activation in our system.

Finally, we assessed the possible involvement of mitochondrial membrane perturbation prior to inflammasome activation and pyroptosis, an event that could also be linked to the activity of ESX-1 and related proteins. Mitochondrial damage has been strongly implicated in inflammasome activation in general, and in Mtb-induced necrosis ^25, 29, 33, 34, 55^. In particular, Mtb H37Rv is reported to cause a loss in mitochondrial membrane potential (ΔΨ_m_), which could be inhibited by cyclosporine A (CsA), a cyclophilin D inhibitor that prevents formation of the mitochondrial permeability transition pore ^34, 56^. However, we did not see an effect of CsA on Mtb-induced inflammasome activation and pyroptosis after 24 hours of infection (Figure 3c). In addition, we monitored the ΔΨ_m_ by live-cell imaging of cells infected with Mtb-BFP and labelled with tetramethylrhodamine ethyl ester (TMRE), a dye that is accumulated in mitochondria in proportion to ΔΨ_m_ ^57^. We observed that ΔΨ_m_ was stable in infected cells until ASC speck formation, and that the potential quickly dropped after ASC speck formation (measured by a drop in TMRE intensity, Figure 3d and Movie S5). This immediate drop in TMRE intensity after ASC speck formation was abolished in THP-1 GSDMD KD macrophages, without any adverse effects on the ability of the cells to form ASC specks, suggesting that GSDMD activity and possibly PM disruption during cell death is required for mitochondrial destabilization (Figure 3e, Movie S6). Similar results were obtained with LPS and nigericin treatment, where ΔΨ_m_ dropped only after ASC speck formation, and the drop was inhibited in GSDMD KD cells (Figure S3b and Movies S7 and S8). We conclude that gross mitochondrial membrane disruption is not a prerequisite for inflammasome activation caused by Mtb or LPS+nigericin but rather accompanies pyroptosis, which is also in line with the intact mitochondrial morphology observed by EM in living cells with an assembled inflammasome (Figure 2c).

### ESX-1-mediated phagosomal damage is a prerequisite for NLRP3 activation by phagosomal Mtb

One of the main effects of the ESX-1 secretion system is destabilization of phagosomal membranes ^36–38^. We therefore asked how phagosomal rupture is related to inflammasome activation and pyroptosis. Galectin-3 (Gal3) binds to exposed glycosylated proteins usually confined to the inner phagosomal membrane and has been used as a marker for ruptured phagosomes ^16, 58^. We infected THP1-Gal3-mScarlet macrophages and observed that Gal3 was indeed recruited to the vicinity of wild-type Mtb, but not to MtbΔRD1 (Figure 4a), and Gal3 recruitment was enhanced upon infection with Mtb grown in the absence of tween (Figure 4b). Gal3 recruitment to Mtb phagosomes occurred prior to inflammasome activation in about 80% of pyroptotic cells, and a typical sequence of events is depicted in Figure 4c and Movie S9.

**Figure 4.**
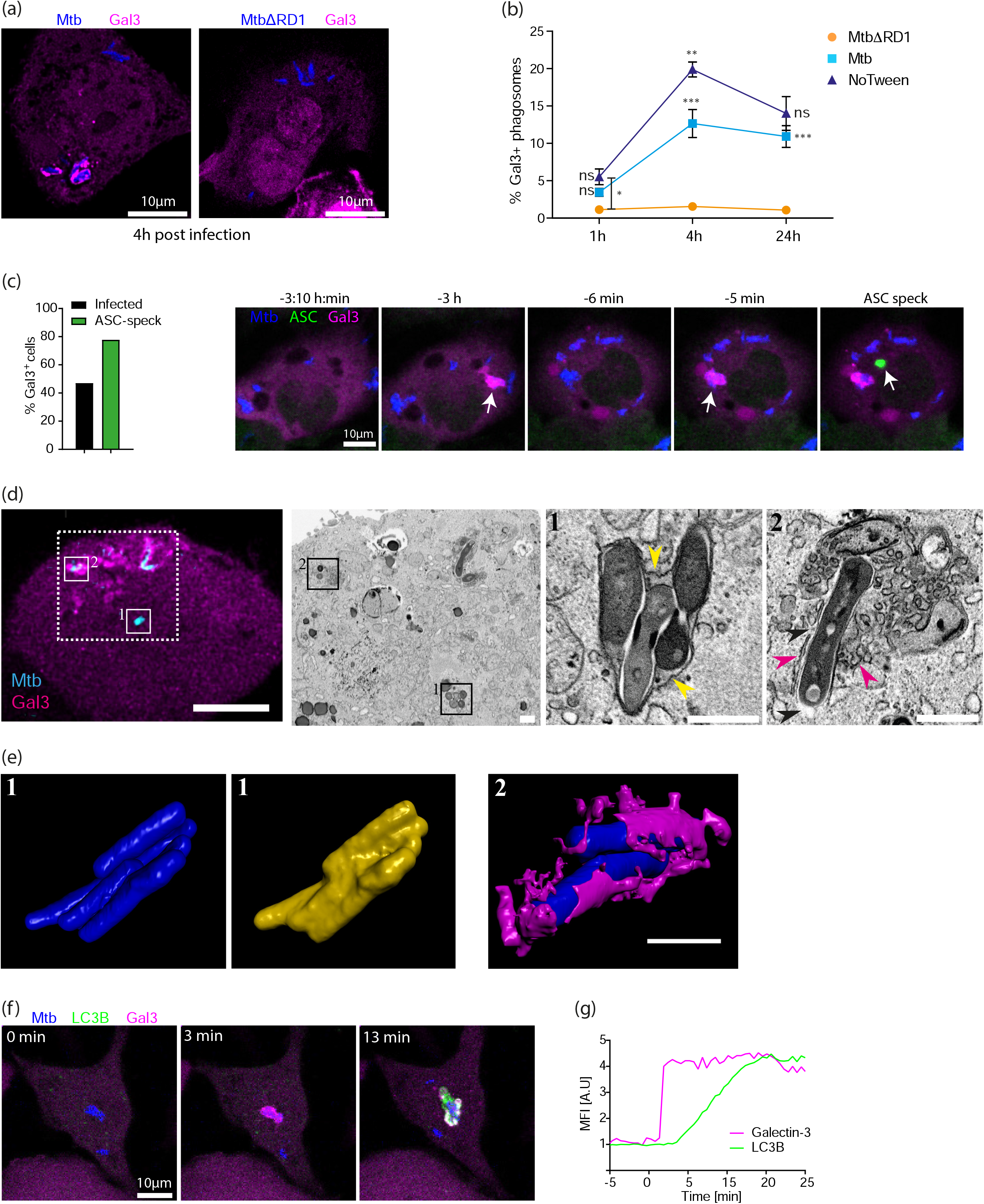
ESX-1-mediated phagosomal damage is a prerequisite for NLRP3 activation by phagosomal Mtb. (**a**) Representative images of THP1 Galectin3-mScarlet (magenta) cells infected with MtbH37Rv::BFP or Mtb H37RvΔRD1::BFP (blue), 4h p.i., (**b**) THP1 Gal3-mScarlet cells were fixed at indicated time points post infection with Mtb H37Rv::BFP, Mtb H37Rv ΔRD1::BFP or Mtb H37Rv::BFP cultured with or without tween-80 (all MOI 20) and imaged by confocal microscopy. The percentage of Gal3^+^ phagosomes was quantified (n>4000 phagosomes per condition, 3 independent experiments) by automated image analysis. Phagosomes with more than one bacterium were considered single phagosomes. *p<0.05, **p<0.01, ***p<0.001, by one-way ANOVA with Tukey’s test. Statistics are calculated compared to the closest lower condition, except for the star between ΔRD1 and noTween at 1h as indicated. Mean±SEM shown. (**c**) THP1 ASC-GFP/Galectin3-mRuby3 cells were imaged by confocal time-lapse for 24h, and the occurrence of Mtb-associated Gal3+-events in infected, surviving cells (n=85) compared to cells undergoing pyroptosis (n=56 cells) was quantified. A representative time-lapse showing Gal3-mRuby3 (magenta) accumulation at Mtb::BFP phagosomes (blue) prior to ASC speck (green) formation. (**d**) Correlative confocal and FIB-SEM imaging of a non-pyroptotic THP1 ASC-GFP/Galectin3-mRuby3 (magenta) macrophage infected with Mtb H37Rv::BFP (cyan) for 24 hours. Dashed square indicates FIB-SEM area, and higher magnification insets are numbered (1) and (2) for Gal3^-^ and Gal3^+^ Mtb. Yellow arrowheads in (1) indicate the intact phagosomal membrane; magenta arrowheads in (2) indicate patches of host cell membrane that are recruited around a bacterium, while absence of membrane is indicated by black arrowheads. Scale bars are 10 µm, 5 µm, 1 µm and 1 µm respectively. (**e**) 3D reconstructions of FIB-SEM tomography data of the bacteria from insets 1 and 2 (blue), along with the corresponding intact phagosomal membrane (yellow) or patches of host cell membrane (magenta). (**f**) THP1 mNeonGreen-LC3B (green)/Galectin3-mScarlet (magenta) macrophages were infected with Mtb H37Rv::BFP (blue) and imaged by time lapse microscopy. (**g**) Normalised fluorescent intensities of one representative Gal3 and LC3B event of n>280 events in 2 independent experiments is shown. *See also Figure S4*.

Using CLEM, we confirmed that Gal3 accumulation corresponds to cytosolic contact of Mtb (Figure 4d). Mtb in phagosomes devoid of Gal3 were surrounded by a tightly apposed and continuous phagosomal membrane, in contrast to bacteria associated with Gal3 which had no visible phagosomal membrane and were in direct contact with the host cell cytosol. Furthermore, the CLEM approach revealed in great detail the presence of clusters of vesicles and membranous structures adjacent to Gal3^+^ bacteria, but not the Gal3^-^ and intact Mtb phagosomes. The ultrastructure is similar to that previously observed for Mtb residing in LC3^+^ compartments in lymphatic endothelial cells ^59^. We therefore investigated the recruitment of LC3B in relation to Gal3 by live-cell imaging of Mtb-BFP-infected THP-1 macrophages expressing mNeonGreen-LC3B and Gal3-mScarlet. Indeed, we consistently observed recruitment of LC3B to intracellular Mtb shortly after recruitment of Gal3 (Figure 4f, g and Movie S10), suggesting that ruptured Mtb phagosomes are targeted by autophagy ^60–62^. To investigate if autophagic targeting of Mtb lead to formation of mature, acidified autophagosomes, THP-1 macrophages expressing Gal3-mScarlet were labelled with LysoView633 (Figure S4a). Only ∼20% of Mtb in ruptured phagosomes were later found in acidified compartments indicating that autophagy of Mtb after phagosomal rupture is inefficient (Figure S4b).

### Inflammasome activation by Mtb is independent of lysosomal damage and release of active cathepsins

The ability to disrupt phagosomes seems important for inflammasome activation by Mtb, and it has been reported that phagosomal acidification is a prerequisite for the membrane damaging activity of ESX-1 ^63–65^. Additionally, active cathepsin release from ruptured phagolysosomes is a suggested trigger for the NLRP3 inflammasome ^24, 39, 40^. We therefore went on to investigate the possible involvement of phagosomal acidification and release of active cathepsins in inflammasome activation and pyroptosis by Mtb. Live-cell imaging of THP1-Gal3-mScarlet or THP1-ASC-GFP cells in the presence of the pH sensitive dye LysoView633 revealed that Mtb was equally efficient at escaping from acidified and neutral compartments (Figure 5a-c and Movies S11 and S12), suggesting that phagosomal acidification is not a prerequisite for Mtb escape into the cytosol. The LysoView signal in single cells was stable prior to ASC speck formation, indicating that there is no general disruption or degradation of acidified lysosomes prior to inflammasome activation (Figure 5d-e and Movie S13). In addition, we did not see any effect on cell death or ASC speck formation when cells were treated with Bafilomycin A1 (BafA1) which inhibits lysosomal acidification by V-ATPase, or the pan cathepsin inhibitor K777, during infection (Figure 5f), despite efficient cathepsin B inhibition (Figure S5). Compared to K777, cathepsin B inhibitor Ca-074-Me was less potent and had clear off-target effects on inflammasome activation during LPS+nigericin treatment (Figure S5). These results demonstrate that Mtb causes inflammasome activation and pyroptosis independently of phagolysosomal acidification and release of active cathepsins.

**Figure 5.**
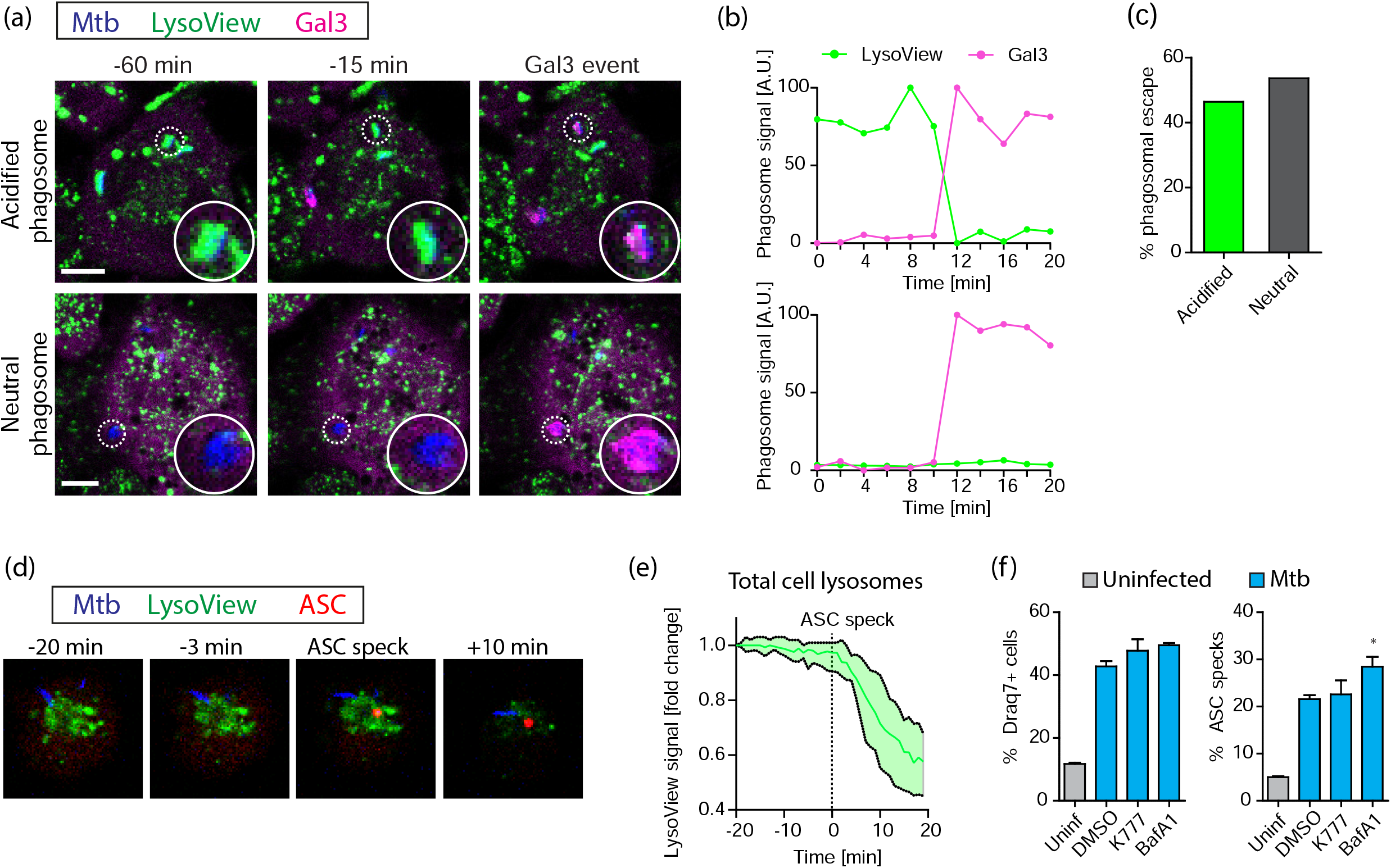
Inflammasome activation by Mtb is independent of lysosomal damage and release of active cathepsins. (**a**) Representative images from 24 hours of time-lapse confocal imaging of THP1 Galectin-3-mScarlet (magenta) macrophages labelled with LysoView633 (green) and infected with Mtb H37Rv::BFP (blue). (**b**) Representative traces of Galectin-3 and LysoView signals for the Mtb phagosomes highlighted in (a). (**c**) Quantification of the occurrence of Mtb-associated Gal3+ events occurring on acidified (LysoView+) and neutral (LysoView-) phagosomes. n=69 Mtb phagosomes were analysed. (**d**) Representative images from confocal time-lapse microscopy of ASC speck formation in THP1 ASC-GFP (red) cells labelled with LysoView633 (green) infected by Mtb H37Rv::BFP (blue). (**e**) Quantification of the average LysoView signal in single cells before and after ASC speck formation. n=27 cells analysed, median±IQR shown. (**f**) THP1 ASC-GFP macrophages were treated with DMSO, K777 (15 µM) or BafA1 (50 nM) and infected with Mtb H37Rv. After 24h, ASC specks and DR+ cells were quantified for n>2000 cells per condition, and the data were analysed by one-way ANOVA with Dunnett’s test. Mean±SEM shown. Data representative of 3 independent experiments. See also Figure S5.

### Mtb carrying ESX-1 can directly damage the host cell plasma membrane

K+ efflux precedes NLRP3 activation for a range of known triggers ^28^. Common for many of those triggers is that they make the PM permeable to K+, e.g. by opening a membrane resident pore (P2X7, pannexin-1) or by inserting and making new pores (MLKL, complement, bacterial pore forming toxins) ^66^. We and others have shown that Mtb-induced NLRP3 inflammasome activation is dependent on K+ efflux, and it has also been shown that Mtb ESX-1 can be haemolytic ^67^. Moreover, PM damage has been observed during Mtb infection and PM repair protects macrophages from necrotic cell death ^68^. However, whether Mtb could directly cause PM damage or if PM damage is a consequence of Mtb induced intrinsic cell death pathways, was not addressed. We therefore wanted to investigate if Mtb could damage the macrophage PM, and if this could be involved in inflammasome activation by Mtb. To monitor PM integrity, we made a THP-1 reporter cell line with an mNeonGreen fluorescent protein tag on the Ca2+ binding protein Apoptosis-Linked Gene 2 (ALG-2), which is recruited to sites of PM damage by calcium influx ^69–71^. We imaged Mtb-BFP infected THP-1 macrophages by time-lapse confocal microscopy and observed ALG-2 recruitment in infected cells, almost 90% of which occurred in close vicinity to Mtb bacteria (Figure 6a, b and Movies S14 and S15). We observed ALG-2 recruitment to Mtb both without prior Gal3 recruitment and subsequent to Gal3 recruitment, with a distribution close to 50/50 between the two categories. This suggests that Mtb can damage the host cell PM either during phagocytosis or following phagocytosis and phagosome rupture.

**Figure 6.**
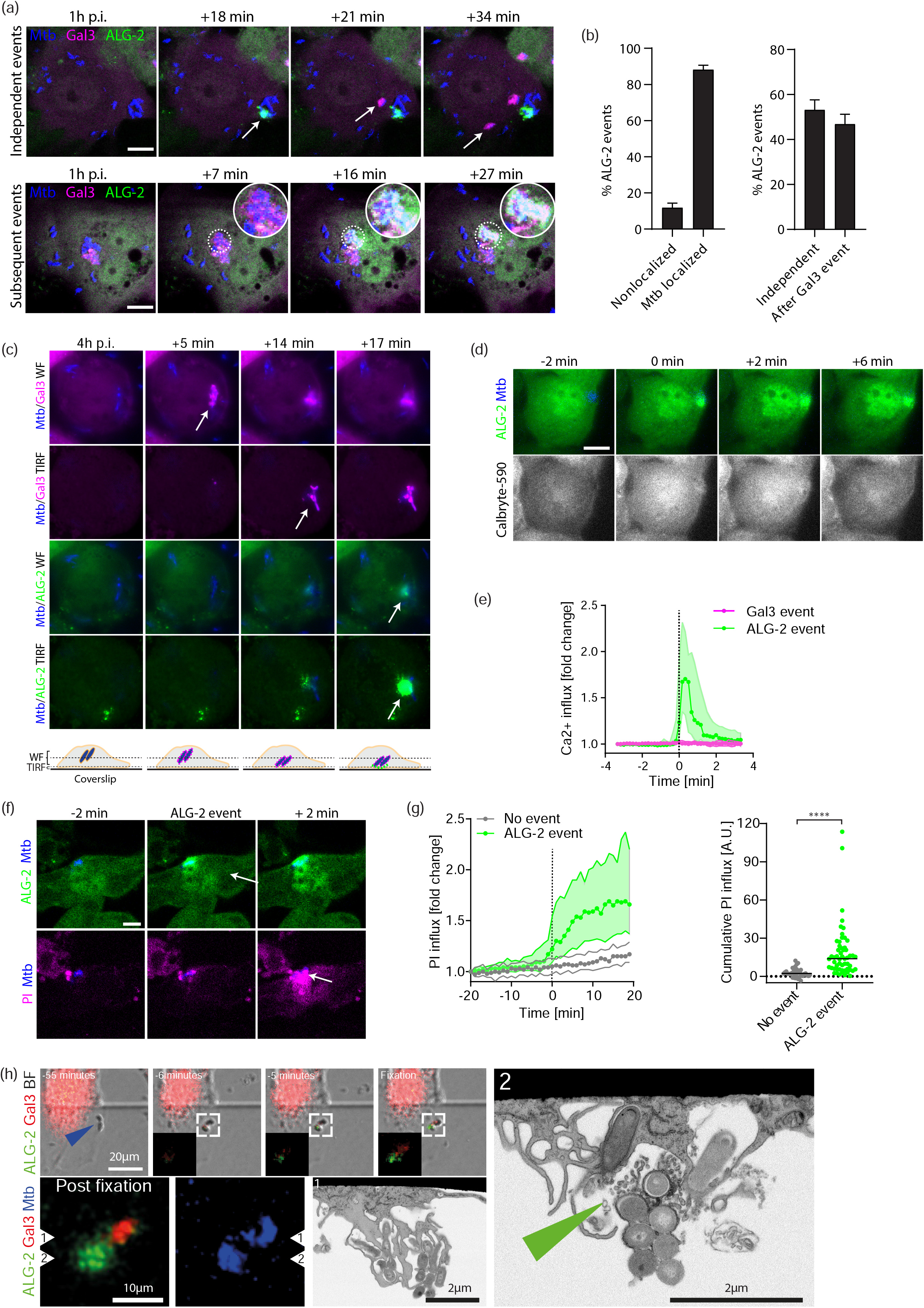
Mtb carrying ESX-1 can directly damage the host cell plasma membrane. THP1 Galectin3-mScarlet/mNeonGreen-ALG-2, THP1 mNeonGreen-ALG-2 or THP1 mNeonGreen-ALG-2/Galectin-3-SNAP(SiR) macrophages were infected with Mtb H37Rv::BFP or Mtb H37Rv mc^2^6206 stained by eFluor450 and imaged live by confocal (Rv::BFP) or widefield and TIRF microscopy (Rv mc^2^6206). (**a**) Representative confocal time-lapse data showing ALG-2 (green) recruitment to plasma membrane-Mtb (blue) contact points either independent of or after Galectin-3 (magenta) recruitment. (**b**) Quantification of ALG-2 events independent of or after Galectin-3 recruitment to Mtb phagosomes and the localization of ALG-2 events. n=213 and n=321 events from 3 independent experiments analysed. (**c**) Simultaneous widefield (WF) and TIRF time-lapse imaging of ALG-2 (green) and Galectin-3 (magenta) during Mtb infection (blue). Representative of n>10 experiments with similar events. (**d**) Time-lapse of Ca^2+^ influx (Calbryte-590, grey) upon ALG-2 (green) recruitment to Mtb (blue). (**e**) Quantification of Calbryte-590 signal during ALG-2 recruitment and Gal3 recruitment events (n=10 events per condition). Median±IQR shown. (**f**) Representative time-lapse images of local extracellular propidium iodide (PI, 20µg/mL) influx during ALG-2 (green) recruitment to Mtb H37Rv::BFP (blue). (**g**) Quantification of PI signal and the cumulative PI influx in single cells during ALG-2 events (n=55) compared with neighbouring cells without ALG-2 events (n=37). Median±IQR and median values shown, representative of 2 independent experiments. ****p<0.0001 by Mann-Whitney test. (**h**) Time-lapse confocal imaging of Mtb-localized ALG-2 event indicated by arrow in THP-1 mNeonGreen-ALG-2 cells infected by Mtb H37Rv::BFP. After time-lapse imaging, cells were fixed and imaged by confocal and FIB-SEM tomography. The FIB-SEM imaging plane is perpendicular to the confocal image and indicated by triangles. Location of ALG-2 on confocal image is indicated by green triangle in FIB-SEM. Confocal scale bars: 10 µm. *See also Figure S6*.

As damage to the PM from the cytosolic side by already internalized bacteria has (to our knowledge) not been described before, we wished to verify that ALG-2 recruitment to Gal3-positive Mtb indeed occurred at the PM. TIRF microscopy is only sensitive to the region within ∼100 nm from the substrate, meaning it specifically detects events occurring at the PM ^72^. We imaged infected THP-1 macrophages with widefield and total internal reflection (TIRF) microscopy simultaneously (Figure 6c and Movie S16). From widefield imaging we could see Gal3 recruitment to intracellular Mtb that were out of range for TIRF, while ALG-2 recruitment was observed in TIRF and WF modes when Mtb came into the range of the TIRF excitation, demonstrating that ALG-2 recruitment following Gal3 also occurs at the PM.

To better understand the extent of PM damage marked by ALG-2 events, we imaged cytosolic calcium levels using the fluorescent indicator Calbryte-590 during time-lapse microscopy of Mtb-infected THP-1 cells. We observed a consistent influx of calcium at timepoints of ALG-2 recruitment (Figure 6d, e and Movie S17), which is expected considering that ALG-2 is known to be recruited by calcium influx. Calcium influx was not observed at timepoints of Gal3 recruitment to Mtb phagosomes (Figure 6e, Figure S6a and Movie S18). We also imaged infected THP-1 macrophages with the membrane impermeable RNA/DNA dye propidium iodide (PI) present in the medium. We noted a consistent local influx of PI in regions of Mtb-localised ALG-2 recruitment events, in contrast to control cells with no apparent ALG-2 events (Figure 6f, g, Movie S19). Taken together, these data demonstrate that Mtb can damage the macrophage PM in a manner depending on close proximity between the two. The damage caused by Mtb makes the PM permeable to both ions (Ca^2+^, K^+^) and larger molecules such as PI.

To investigate the ultrastructure of Mtb-associated PM damage we performed live-cell CLEM. After live-cell imaging and rapid fixation, recent ALG-2 recruitment events were imaged by confocal or Airyscan microscopy, and further by FIB-SEM tomography (Figure 6h, Figure S6b). At the location of fluorescent ALG-2 signal and directly adjacent to Mtb, multiple 50-100 nm sized vesicles were visible, indicative of ESCRT-mediated PM repair ^70, 73^.

### Plasma membrane damage caused by Mtb or Silica activates the NLRP3 inflammasome

Knowing the dependency on K+ efflux for NLRP3 activation, we hypothesized that the ability of Mtb to cause PM damage (hence making it permeable to ions) would be linked to inflammasome activation. We therefore imaged THP1-ALG2-mNeonGreen/Gal3-mScarlet/ASC-mIRFP670 macrophages by time-lapse microscopy during infection with Mtb-BFP. An example time course is shown in Figure 7a and Movie S20. We recorded the time points of all Gal3 (phagosomal rupture) and ALG-2 (PM damage) recruitment events in cells forming ASC specks, along with the time point of ASC speck formation. PM damage preceded ASC speck formation in 87% of cells infected with Mtb and occurred closer in time to ASC speck formation than Gal3 recruitment to Mtb phagosomes (Figure 7b). While the median time from phagosomal rupture to ASC speck formation was 2 h, the median time from PM damage to ASC speck formation was 1 h. Moreover, within 20 minutes before ASC speck formation, 48% of cells had ALG-2 events and 23% of cells had Gal3 events, compared to 1% and 4%, respectively, in infected control cells over an average 20-minute period (Figure 7c). Thus, especially PM damage events are strongly enriched in the time before ASC specks, connecting these events at the single-cell level. When cells were infected with MtbΔRD1, which does not damage host cell membranes, we did not observe any ALG-2 recruitment and ASC speck formation was significantly reduced, as noted earlier (Figure 3a). Treatment with the phagocytosis inhibitor cytochalasin D during infection with Mtb only partially inhibited ASC speck formation after 24 hours, supporting that Mtb can induce inflammasome activation through PM damage from outside of the cell (Figure 7d). Finally, when inflammasome activation was inhibited by MCC950 or extracellular KCl during infection, the number of ALG-2 events per cell remained constant while ASC speck formation was inhibited as earlier, confirming that PM damage happens prior to NLRP3 inflammasome activation (and is not caused by e.g. active GSDMD) (Figure 7e and Figure S7a).

**Figure 7.**
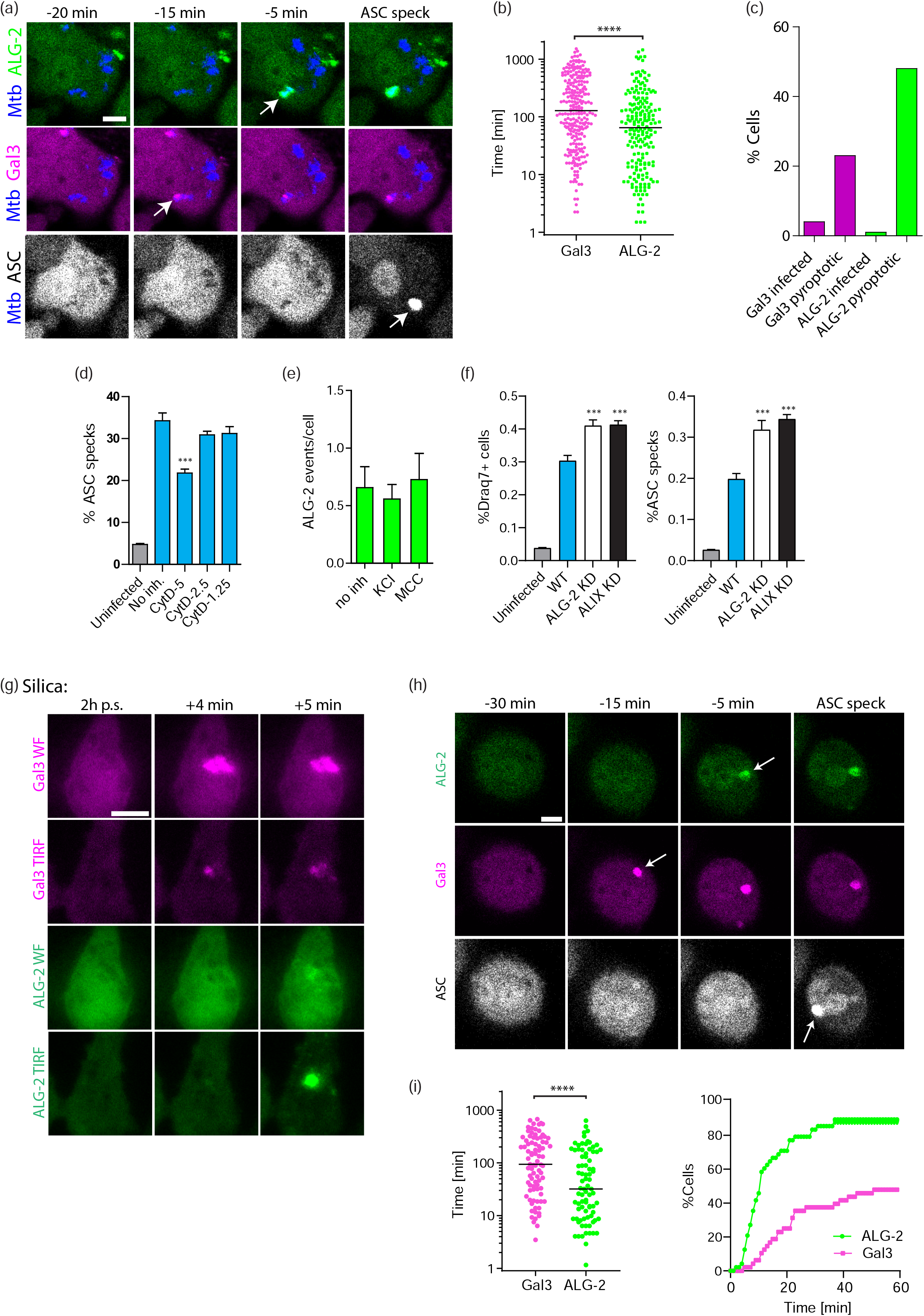
Plasma membrane damage caused by Mtb or Silica activates the NLRP3 inflammasome. THP1 mNeonGreen-ALG-2/Galectin-3-mScarlet/ASC-mIRFP670 cells were infected by Mtb H37Rv::BFP (MOI 10) and imaged by time-lapse microscopy for 24h. (**a**) Representative time-lapse of ALG-2 (green), Gal3 (magenta) and ASC (grey) dynamics during Mtb infection (blue). (**b**) Timing of Mtb-localized Gal-3 and ALG-2 events compared to ASC speck formation in all cells forming ASC specks. n=200 ALG-2 and n=262 Gal-3 events analysed from 3 independent 24h time-lapse experiments. Lines indicate median values. ****p<0.0001 by Mann-Whitney test. (**c**) Percentage of cells with ALG-2 and Gal3 events in the population of cells forming ASC specks (pyroptotic) within 20 minutes before ASC speck formation, compared to the percentage of cells in the surviving infected population with events in an average 20 min period. (**d**) % ASC specks in cells infected with Mtb H37Rv::BFP MOI5 for 24h in the presence or absence of the indicated concentration (in µM) of the inhibitor cytochalasin D (**e**) Average number of Mtb-localized ALG-2 events per cell during 24h infection in the absence or presence of KCl (40mM) or MCC950 (10µM). (**f**) Quantification of DR+ cells and ASC specks 24h p.i. (MOI 20) in THP1 ASC-mNeonGreen (WT) cells and WT cells depleted of ALG-2 or ALIX by CRISPR-Cas9. n>2000 cells per condition, and the data was analysed by one-way ANOVA with Dunnett’s test. Mean±SEM shown, *p<0.05, **p<0.01. Data representative of 2 independent experiments. (**g**) Simultaneous widefield (WF) and TIRF time-lapse imaging of ALG-2 (green) and Gal-3 (magenta) during 4h treatment with silica (100µg/mL). (**h**) Representative confocal time-lapse images of a cell forming an ASC speck (grey) during 24h treatment with 100µg/mL silica. (**i**) Cells were treated with 100µg/mL silica. Timing of Gal-3 and ALG-2 events compared to ASC speck formation in all cells forming ASC specks. n=82 ALG-2 and n=85 Galectin-3 events analysed from 3 independent 24h time-lapse experiments. Lines indicate median values. Quantification of the percentage of cells with ALG-2 or Gal-3 events in the given time interval before ASC speck formation in single cells. n=48 cells from 3 independent 24h time-lapse experiments. ****p<0.0001 by Mann-Whitney test. Confocal scale bars: 10 µm. *See also Figure S7*.

ALG-2 and the ALG-2 interacting protein X (ALIX) are early components of ESCRT-mediated PM repair ^69, 70^. We went on to inhibit PM repair by knocking down ALG-2 or ALIX by CRISPR-Cas9 in THP-1 macrophages. Mtb caused significantly more inflammasome activation and pyroptosis both in ALG-2 and ALIX KD macrophages after 24 hours of infection, compared to control cells (Figure 7f and Figure S7b). Together, these results demonstrate that Mtb-induced PM damage is a trigger for NLRP3 inflammasome activation in macrophages, and that PM repair mechanisms negatively regulate inflammasome activation and pyroptosis caused by Mtb.

Silica is another NLRP3 stimulus that supposedly triggers inflammasome activation by destabilizing host cell phagolysosomes after uptake ^24^. We therefore thought it interesting to study NLRP3 activation by silica in the same system as we used for Mtb infection. TIRF and widefield microscopy of Gal3 and ALG-2 tagged THP-1 macrophages revealed Gal3 recruitment to intracellular silica crystals and ALG-2 recruitment to sites at the PM (Figure 7g and Movie S21), indicating that silica crystals are also capable of disrupting phagosomal membranes and the PM. A typical time course of events culminating in ASC speck formation is shown in Figure 7h and Movie S22. PM damage preceded ASC speck formation after silica stimulation in most cells (90%). Further, silica-induced ALG-2 events were closer in time to ASC specks than Gal3 events, similar to what we observed during Mtb infection (Figure 7i). 71% of cells that assembled ASC specks had an ALG-2 recruitment event within the last 20 minutes before ASC speck formation, while about 25% of the cells had a Gal3 recruitment event within the same period (Figure 7i). Nigericin, which directly depletes the cell of K^+^ as a K^+^ ionophore, or imiquimod, which is an NLRP3 trigger acting independently of K+ efflux ^29^, did not result in any visible Gal3 or ALG-2 recruitment events prior to inflammasome activation (FigS7c, d and Movies S23 and S24).

Further, we investigated if lysosomal damage using the lysosome-targeted photosensitizer TPCS_2a_ ^74^ could activate NLRP3 (Figure S7e, f and Movie S25). However, despite loss of lysosomal pH and Gal3 recruitment indicating lysosomal damage after blue light excitation in the presence of TPCS_2a_, we did not observe ALG-2 events at the PM or ASC speck formation. These data show evidence that PM damage (with subsequent K+ efflux) is a central event upstream of NLRP3 inflammasome activation by the “lysosomal triggers” Mtb and silica, and that lysosomal damage itself is not sufficient to activate NLRP3.

## Discussion

Studies of the molecular mechanisms involved in the interaction between *Mycobacterium tuberculosis* and host cells are complicated by the variability of this interaction. Bacterial containment, replication and host cell death through different programmed or unregulated death pathways are all possible outcomes. The heterogeneity makes average measures hard to interpret, and they are unsuited to distinguish between cause and effect. Long-term studies at the single cell level give a better insight into these questions, allowing to study separate populations in a cell culture and multiple events in the same cell and over time. We capitalize on this advantage to gain several insights into Mtb-host interactions. We focus on the temporal regulatory events that orchestrate NLRP3 activation and pyroptosis, two phenomena that are still not well understood, especially in the context of a complex pathogen infection. Both AIM2 and NLRP3 have been implicated in IL-1β release during Mtb infection ^15, 16, 18, 19^, while the occurrence of pyroptosis as a distinct route of cell death has been less clear ^75, 76^. Here we establish that NLRP3 is activated and that caspase1- and GSDMD-dependent pyroptosis is an important cell death mechanism during Mtb infection in human cells, together leading to release of IL-1β. Intriguingly, NLRP3 activation and pyroptosis is triggered by PM damage caused by Mtb during phagocytosis or after phagosomal escape. Similar results were obtained with silica crystals, suggesting PM damage as a common mechanism of NLRP3 activation and pyroptosis by sterile and pathogenic agents holding membrane disruptive properties.

Necroptosis is another necrotic cell death pathway induced by Mtb, suggested to be caused by the secreted enzyme TNT ^33, 34^. In some cases, there is cross-talk between necroptosis and NLRP3 activation and pyroptosis ^77, 78^, but our results indicate that these are distinct processes during Mtb infection. We do not see an effect of necroptosis inhibitors or TNT mutants, and by time-lapse imaging we show that destabilization of mitochondria accompanies pyroptosis rather than causes inflammasome activation in the NLRP3 pathway. The mitochondrial integrity is similar both in response to Mtb and to LPS+nigericin. The discrepancy between our results and those of Pajuelo et al. ^34^ with regards to cell death by TNT and necroptosis inhibitors is likely due to the longer infection time used by Pajuelo and colleagues: our analyses are mostly performed after 24 hours of infection where pyroptosis is most prominent, but we also see an increase in necrotic cell death at later timepoints.

At the single cell level, highly infected cells at earlier stages of infection (first 24h) are the most prone to pyroptosis, and there is a strong correlation between ESX-1 activity damaging host cell membranes and NLRP3 activation. Other groups have also shown the correlation between bacterial burden and cell death ^76, 79, 80^. One could hypothesize that *in vivo* pyroptosis would occur during a phase of infection with presence of larger amounts of Mtb, such as during active phases of the disease where Mtb needs to spread ^80^. Interestingly, although IL-1β is a central cytokine in a successful immune response against tuberculosis ^81^, a gain of function mutation in NLRP3 has been associated with poor clinical outcome in tuberculosis patients ^82^.

The precise mechanism of NLRP3 activation has evaded the community for over a decade. Although there are alternative routes proposed, inhibition of NLRP3 activation by high concentrations of extracellular KCl positions K+ efflux as the predominant upstream effector of NLRP3 activation by a wide range of triggers. This is also the case for Mtb, as we and others have shown ^14, 39^. The similarities in ASC speck structure and progression of pyroptosis during Mtb infection and the canonical NLRP3 trigger nigericin, which directly depletes the cell of K+ as a K+/H+ ionophore, indicates that potential differences are mainly linked to how K+ efflux occurs. Lysosomal damaging agents are one of the most clinically relevant classes of NLRP3 triggers ^83^. Cathepsin release from damaged lysosomes, especially of Cathepsin B, is one commonly proposed connection, although how this further causes K+ efflux has been unclear ^24, 39, 40^. One main source of this confusion is the widespread use of the Cathepsin B inhibitor Ca-074-Me, which appears to inhibit NLRP3 activation independent of Cathepsin B inhibition (our results and ^28, 41–43)^. We also show that although Mtb can damage acidified phagolysosomes, this is not required for NLRP3 activation, and there is no general loss of lysosomal content prior to NLRP3 inflammasome activation. Instead, PM damage appears to be the key event to trigger NLRP3 during Mtb infection. Also, in response to silica crystals, a canonical lysosome-damaging NLRP3 trigger, we observe PM damage prior to NLRP3 activation. Permeabilization of PM by e.g. bacterial pore-forming toxins is well known to cause ion fluxes down their electrochemical gradients and thus activate NLRP3 through the K+ efflux route ^66^. PM damage during phagocytosis of Mtb, silica and likely other similar membrane damaging agents explains some of the NLRP3 activation events, but there is also a second possible route. Our imaging data show that PM damage can also occur from the cytosolic side after uptake, if Mtb or silica has damaged the phagosome and subsequently come into contact with the PM. Thus, a straight forward explanation for how something that primarily damages phagolysosomes can cause K+ efflux and NLRP3 activation emerges. Strikingly, the opposite case with extensive, but localized and controlled lysosomal damage caused by a lysosomal photosensitiser does not cause NLRP3 activation. Since PM damage is actively repaired through many routes, including through Ca2+ - dependent ESCRT machinery ^44, 69–71^, it is also interesting to note that this branch of ESCRT emerges as a negative regulator of NLRP3 activation and cell death during Mtb infection. The ESCRT machinery was also recently shown to be involved in repair of phagosomes damaged by mycobacteria ^84, 85^ and to regulate pyroptosis or necroptosis by repairing GSDMD pores ^86^ or MLKL pores ^87^, respectively, further pointing to the importance of the ESCRT machinery in regulation of inflammation and securing cell viability.

How ESX-1 damages membranes is not well understood. Several papers indicate a central role for the secreted protein ESAT-6, acting through membrane lysis or membrane pore formation ^35, 63, 65, 88, 89^. However, this has recently been attributed to detergent contamination, and a contact-induced form of gross membrane disruption caused by undefined ESX-1 activity has been proposed ^67^. ESX-1 secreted substrates are also retained in the mycobacterial capsule, and ESAT-6 has been observed to quickly disappear upon contact between bacteria and host macrophages ^51, 52^. Our results also indicate contact-induced damage being most likely, as damage events are localized to contact points between Mtb and the PM. The damage upon contact could be caused by ESX-1-dependent factors in the bacterial capsule or other factors that are diminished by the presence of the detergent tween, as Mtb cultured without tween showed increased membrane damaging activity and higher inflammasome activation and cell death. The damage also appears to be severe, as shown by ultrastructural investigations, the simultaneous recruitment of Gal3 and loss of lysosomal pH in phagolysosomes, and the simultaneous recruitment of ALG-2 and PI and calcium influx during PM damage. This is different from previously described bacterial toxins that cause membrane damage by pore formation. Furthermore, the membrane damaging activity of ESX-1 is unique in that it can act from both sides of the PM. Our ultrastructural investigations of sites of PM damage close to Mtb indicate that the PM repair response in the affected cell is quickly initiated, complicating a direct observation of what the damage initially looks like.

In conclusion we show a K+ efflux dependent, NLRP3-caspase-1-GSDMD axis for pyroptotic cell death and IL-1β release during Mtb infection of human macrophages and monocytes. We emphasize the importance of membrane damage through activity of the ESX-1 secretion system, upstream of NLRP3 activation at the single cell level. Rather than phagolysosomal damage itself causing NLRP3 activation, we show that PM damage occurring during phagocytosis or from the cytosol after phagosomal damage is the cause of K+ efflux and subsequent NLRP3 activation by Mtb. In this scenario, NLRP3 acts as a sensor for cell integrity rather than as a sensor for Mtb in the cytosol or release of lysosomal enzymes.

Mtb needs to escape from host macrophages to efficiently replicate and spread to new hosts. Based on our data we propose a model where preservation of membrane integrity is key to contain Mtb infections. The ESCRT machinery repairs damages caused by Mtb to the plasma membrane and phagosomes whereas autophagy sequesters ruptured Mtb phagosomes. Inefficient countermeasures by the host cell allows for potassium efflux and NLRP3 inflammasome activation, in which case pyroptosis is inevitable and facilitates the spread of Mtb to neighbouring cells and eventually to new hosts. Cell necrosis pathways are thus attractive as possible targets for host-directed therapeutic strategies ^90^. We also show that PM damage is a shared mechanism with the crystalline NLRP3 trigger silica. Crystalline triggers cause sterile inflammatory diseases in the lungs and the cardiovascular system, and the mechanism of NLRP3 activation by these so-called lysosomal triggers has so far been elusive. Our findings point to PM damage as a shared mechanism for this group of NLRP3 activators, extending the relevance beyond tuberculosis disease.

## Acknowledgements

We thank Liv Ryan and Unni Nonstad for FACS of THP-1 cells, Katherine Fitzgerald and Egil Lien at UMass for reading and commenting on the manuscript, and members of the mycobacterial- and HIV research group at CEMIR for valuable discussions. The fluorescence microscopy work was performed at the Cellular and Molecular Imaging Core Facility (CMIC), NTNU. CMIC is funded by the Faculty of Medicine at NTNU and Central Norway Regional Health Authority. Funding sources: The Research Council of Norway for support to the Norwegian Micro- and Nano-Fabrication Facility (NorFab) project number 245963/F5; the Centre of Excellence grant to CEMIR project number 223255 and FRIPRO project numbers 231303 and 287696 (to THF). NTNU Enabling Technologies for PhD funding (SU).

## Author contributions

Conceptualization: THF, KSB and MSB; Investigation: KSM, MSB, SU, RS, HK, AM, SEÅ, TÅS; Writing – initial draft: THF, KSB, MSB; Writing – review and editing: THF, KSB, MSB; Supervision and Funding acquisition: THF. All authors discussed the results and commented on the manuscript.

## Materials and Methods

All reagents used are listed in supplementary table 1.

### Experimental models

THP-1 cells were maintained in RPMI 1640 supplemented with L-glutamine, 10 mM HEPES and 10% FCS, and passaged regularly to keep the cell density between 0.2 and 1×10^6^/mL. Cell lines were routinely tested for mycoplasma. Human peripheral blood mononuclear cells (PBMCs) were isolated with Lymphoprep from buffy coats obtained from healthy volunteers (both male and female) at the blood bank of St Olavs Hospital (Trondheim, Norway). Collection of human blood was approved by the Regional Committee for Medical and Health Research Ethics in Central Norway.

Mtb strains were grown at 37°C in Middlebrook 7H9 medium supplemented with 0.2% glycerol, 0.05% Tween-80, and oleic acid, albumin, dextrose and catalase (OADC). For Mtb mc^2^6206 auxotroph strain 50µg/mL L-leucine and 24µg/mL D-pantethonate were added to the growth medium.

Lentiviral production was performed in HEK293T cells as described ^91^ using 3^rd^ generation packaging system with pMDL.g/pRRE, pMD2.G and pRSV-Rev and JetPrime (PolyPlus) as transfection reagent. Virus-containing supernatant were harvested 2 and 3 days post transfection and THP-1 cells were transduced in the supernatant by 90 minutes spinoculation at 32°C, 1000g in the presence of 8µg/mL polybrene, and selected in 1µg/mL puromycin, 10µg/mL blasticidine or 100µg/mL Hygromycin for 1 week. Finally, the cells were selected for a consistent moderate expression level of fluorescent constructs by 1-2 rounds of FACS (BD Aria II). CRISPR modified THP-1 cells were used directly as a polyclonal population after puromycin selection. Mtb H37Rv, Mtb H37RvΔRD1 and Mtb mc^2^6206 were transformed with msp12::EBFP2 and selected on hygromycin (65 µg/mL) 7H10 plates (Difco/Becton Dickinson).

## Method details

### Generation of new vectors

Gateway cloning was used to generate lentiviral expression vectors. Human Galectin-3, human ALG-2 (THP-1 cDNA), human ASC, human LC3B (synthetic GeneArt Strings, ThermoFisher), mScarlet (synthetic), mRuby3, mNeonGreen, mIRFP670 and SNAP_f_ tag were PCR amplified and cloned into Gateway pEntry vectors. All pEntry constructs were verified by Sanger sequencing (GATC Biotech) before further use. PuroR in pLex307 was replaced with BlastR or HygR to generate Gateway lentiviral destination vectors with corresponding antibiotic resistances. 2-fragment Gateway recombination with pLex307, pLex307-Blast or pLex307-Hyg were performed to generate ASC-mNeonGreen, ASC-mIRFP670, Galectin-3-mScarlet, Galectin-3-SNAP, mNeonGreen-LC3B and mNeonGreen-ALG-2 constructs. Guide RNAs targeting GSDMD, NLRP3, ALG-2 or ALIX were cloned into LentiCrispr v2 by BsmBI digestion and ligation.

### Verification of CRISPR knock-down

Knock-down in the cell pool was verified by cleavage assay (GeneArt genomic cleavage assay, ThermoFisher), sequencing and TIDE analysis ^92^ and/or western blotting. Antibodies and dilutions for western blot were NLRP3 (1:1000), GSDMD (1:1000), ALG-2 (1:500) and ALIX (1:1000).

### Macrophage and monocyte infection

Before use, THP-1 cells were differentiated at a concentration of 300.000 cells/mL in medium containing 100 ng/mL PMA for 3 days, washed in cell medium and rested 1 day prior to experiments. Primary monocytes were selected by plastic adherence of PBMCs for 1 hour, followed by 3x washing in HBSS, and cultured in RPMI with 10% A+ serum (blood bank of St Olavs Hospital, Trondheim, Norway). Mtb bacteria were grown until OD_600_ was 0.4-0.55 (log phase) then pelleted at 2400g for 10 minutes, resuspended in RPMI with 10% A+ serum to opsonise bacteria prior to infection, sonicated 2 times for 5 s at 70% power (Branson Digital Sonifier, S-450D), and clumped bacteria were removed by centrifuging at 300g for 4 minutes. The supernatant containing bacteria was diluted in RPMI with 10% A+ serum to the indicated MOI assuming 1 OD_600_=3×10^8^ bacteria/mL, and applied to cells for 45 minutes with or without inhibitors, followed by washing in HBSS and replacement of the media to normal cell medium with 10% A+ with or without inhibitors, or Leibovitz L-15 CO_2_-independent medium with 10% A+ for live cell imaging. The typical condition of MOI 20 infection gave about 50% infection rate with 1-20 bacteria per macrophage. Inhibitor concentrations were DMSO control (1:400), zVAD-FMK (50 µM) VX765 (50µM), MCC950 (10 µM), KCl (40mM), Nec1s (10 µM), GSK-872 (5µM), Cyclosporin A (5 µM), K777 (15µM) or Bafilomycin A1 (50 nM). Supernatants were harvested for ELISA or LDH assays after 24 hours, and analysed by human IL-1β kit or LDH cytotoxicity kit according to the manufacturer’s instructions.

### Live/fixed cell imaging

THP-1 cells were seeded in 35mm glass bottom dishes (Ibidi) or 96-well glass bottom plates (Cellvis) as above. For some experiments bacteria were added at a lower MOI and not washed to be able to follow the entire course of infection. Cells were imaged on a Leica SP8 confocal microscope with a 37°C incubator using a 40×1.3 or 20×0.75 oil immersion objective or 10×0.4 air objective at the timepoints indicated in the text. TIRF/Widefield imaging was done on a Zeiss Laser TIRF 3, with a 63×1.46 oil objective and Hamamatsu EMX2 EMCCD camera. For time-lapse imaging by confocal, typically 6 fields of view comprising >500 cells in total were imaged at 30-45 s intervals, while TIRF/WF was done on one to four fields of view comprising 10-40 cells at 10-30s intervals. For some experiments, DRAQ7 (0.15µM), 10 nM TMRE (10 nM), LysoView 633 (1:10 000) or Calbryte-590 (5µM) was preincubated for 30-60 minutes, and added to the cell medium during Mtb infection. Calbryte-590 was only preincubated and not re-added. In cells with SNAP tag, cells were labelled with 1:200 SNAP-Cell 647-SiR for 15 minutes, washed 3x in cell medium and rested for 30 minutes before use. For lysosomal damage by photosensitizer, TPSC_2a_ was added to medium (0.4µg/mL) for 16 h and cells were washed and incubated for 4h without TPSC_2a_. THP1 cells were stained or not stained with Cresyl Violet (1µM, 5 minutes) exposed to 405 nm light (10%, 1 second), and imaged. Galectin-3 accumulation on phagosomes during infection with different Mtb strains was determined by fixing cells in 4% PFA for 20 minutes at 1, 4 and 24 h post infection, and imaging 16 fields of view with 40x objective per well.

### FIB/SEM sample preparation and correlative imaging

For correlative imaging experiments two growth substrates were used, with different optical microscopy and fixation approaches. In the first approach THP1 ASC-GFP Gal3-RFP cells were seeded and infected as above on moulded aclar slides. Aclar substrates were placed in 24-well plates and imaged live using 10x objective as above. After 24 hours, cells were fixed in 2.5% glutaraldehyde in 100mM PIPES for 1h, placed upside-down in 35mm glass bottom dishes and cells of interest were re-imaged using 63×1.2W objective on a Leica SP8 confocal microscope. In the second approach THP-1 cells were seeded and infected as above on moulded polymer coverslips (ibidi) mounted in 8 well sticky-Slides (ibidi). Samples were imaged using 40×1.3oil objective on a Leica SP8 confocal microscope and fixated by adding 8% paraformaldehyde (PFA) and 0.2% GA in 200mM PIPES directly into imaging media at a 1:1 ratio. After 20 minutes fixation media was changed to fresh 4% PFA and 0.1% GA in 100mM PIPES. Cells with interesting events during live microscopy were then re-imaged with a 40×1.2Imm AutoCorr objective on a Zeiss LSM880 microscope using the Airyscan mode. After re-imaging fixation medium was changed to 2% PFA and 2.5% GA in 100mM PIPES and fixated at 4°C over night. Cells were then prepared for FIB/SEM microscopy and relocated as previously described (Beckwith et al. 2015). Briefly, cells were post-fixed and contrasted in potassium ferrocyanide and osmium tetroxide, and further *en block* stained in uranyl acetate. Cells were gradually dehydrated and embedded in Durcupan (Sigma-Aldrich) and cured at 60 degrees for a total of 3 days. Samples were attached to SEM sample stubs with a drop of Durcupan before the last day of curing. Before FIB/SEM imaging, a layer of 40nm platinum/palladium was deposited on the samples. FIB/SEM imaging and tomography experiments were performed on a Helios G4 dual-beam FIB/SEM from FEI, using an acceleration voltage of 2kV or 3kV for the electron beam, immersion mode and the in-column mirror detector. Slice thickness was typically set to 20nm or 25nm. Initial alignments, scaling and simple adjustments (brightness/contrast) were done in Fiji, while 3D reconstructions were performed semi-manually in Avizo or Dragonfly.

### Cathepsin B activity assay

THP1 cells were differentiated in 24-well plates as above. 1 h prior to lysis, inhibitors (K777 and Ca074-Me) were added at concentrations indicated in the text. Cathepsin B activity assay was performed as described ^93^ with some modifications. Briefly, cells were lysed in 200µL of lysis buffer (0.1% Triton X-100, 250 mM sucrose, 20 mM HEPES, 10 mM KCl, 1.5 mM MgCl_2_, 1 mM EDTA, 1 mM EGTA, 0.5 mM Pefabloc SC, pH=7.5) on ice for 20 minutes. 50µL of lysate was mixed with 50µL of Cathepsin B reaction buffer (50 mM sodium acetate, 4 mM EDTA, 0.5 mM Pefabloc SC, 8 mM Dithiothreitol (DTT) and 50 µM Z-RR-AMC, pH=6.0), incubated at 30°C for 5 minutes, and fluorescence intensity was monitored every 30 s for 20 minutes (ex 355/em 460 nm, POLARstar Omega microplate reader).

## Quantification and statistical analysis

### Image analysis

For quantification of cell death and ASC specks at defined time points with and without inhibitors or knock-down, 16 fields of view containing n>2000 cells in total were imaged per condition. CellProfiler was used to automatically segment and count live cells and ASC specks based on the ASC-GFP signal, and dead cells based on the DRAQ7 signal. Bacterial burden per cell was measured from live cell time-lapse movies by mean EBFP2 intensity in infected cells immediately prior to ASC speck formation, and compared to cells surviving the duration of the experiment. The time point for the control cells was chosen to be the average time point for ASC-speck induction. Traces of DRAQ-7, TMRE or LysoView signal were generated by manually selecting ROIs around cells with or without ASC specks forming.

Galectin-3 association with Mtb was scored by a trained model in Ilastik and further analysed by CellProfiler. During live-cell imaging the time point for galectin-3 or ALG-2 recruitment events was determined by the first visible accumulation at a bacterial phagosome in a cell later forming an ASC speck. Data were plotted and statistically analysed using Python or GraphPad Prism, while images and figures were prepared using FIJI and Adobe Illustrator. TIRF/WF images of mNeonGreen-ALG-2 and Galectin-3 mScarlet were unmixed by linear unmixing due to substantial cross-excitation of mScarlet with 488 nm laser.

### Statistical analysis

Detailed statistical analysis for individual experiments are listed in each figure legend. This includes the statistical test performed, the parameters shown, and number of cells, replicates and independent experiments as appropriate. Data comparing mean values of technical replicates of representative experiments or independent experiments were analysed by two-tailed unpaired Student’s t test (two groups) or one-way ANOVA (more groups). For one-way ANOVA corrections for multiple comparisons were applied for comparisons with one control group (Dunnett’s) or all groups (Tukey’s). For comparisons of non-parametric data (fluorescent intensity or event timing) Mann-Whitney U-test was used instead. GraphPad Prism 8.0 was used to perform all statistical analysis and determine p-values, with p-value<0.05 considered significant.

### Data and Software Availability

The data that support the findings of this study and the custom scripts and pipelines for image analysis are available from the Lead Contact upon reasonable request.

## Supplemental Information

Supplemental Table 1 and Figures 1-7 with legends are attached following the main Figures in the merged manuscript and uploaded as a separate file.

Supplemental Movies 1-25 show time-lapse imaging results underlying main and supplemental Figures and are uploaded to the Immunity server:

**Movie S1.** Confocal time-lapse imaging of THP1 ASC-GFP (green) infected by Mtb::BFP (blue) undergoing pyroptosis in the presence of Draq7 (magenta).

**Movie S2.** Confocal time-lapse imaging of THP1 ASC-GFP (green) infected by Mtb::BFP (blue) undergoing non-pyroptotic necrosis in the presence of Draq7 (magenta).

**Movie S3.** Confocal time-lapse imaging of THP1 ASC-GFP (green) not infected by Mtb::BFP (blue) undergoing apoptosis in the presence of Draq7 (magenta).

**Movie S4.** Confocal time-lapse imaging of THP1 ASC-GFP (green) infected by Mtb::BFP (blue) undergoing ASC speck formation, pyroptosis and bacterial dissemination, in the presence of Draq7 (magenta).

**Movie S5.** Confocal time-lapse imaging of THP1 ASC-mNeonGreen stained with the mitochondrial membrane potential indicator TMRE (red, 10nM) and infected by Mtb::BFP (blue).

**Movie S6.** Confocal time-lapse imaging of THP1 ASC-mNeonGreen (green) depleted for GSDMD by CRISPR, stained with the mitochondrial membrane potential indicator TMRE (red, 10nM) and infected by Mtb::BFP (blue).

**Movie S7.** Confocal time-lapse imaging of THP1 ASC-mNeonGreen (green) stained with the mitochondrial membrane potential indicator TMRE (red, 10nM) and treated with LPS+nigericin.

**Movie S8.** Confocal time-lapse imaging of THP1 ASC-mNeonGreen (green) depleted for GSDMD by CRISPR, stained with the mitochondrial membrane potential indicator TMRE (red, 10nM) and treated with LPS+nigericin.

**Movie S9.** Confocal time-lapse imaging of THP1 ASC-GFP (green)/Gal3-mRuby3 (magenta) infected by Mtb::BFP (blue).

**Movie S10.** Confocal time-lapse imaging of THP1 mNeonGreen-LC3B (green)/Gal3-mScarlet (magenta) infected by Mtb::BFP (blue).

**Movie S11.** Confocal time-lapse imaging of THP1 Gal3-mScarlet (magenta) stained by LysoView633 (green) and infected by Mtb::BFP (blue).

**Movie S12.** Confocal time-lapse imaging of THP1 Gal3-mScarlet (magenta) stained by LysoView633 (green) and infected by Mtb::BFP (blue).

**Movie S13.** Confocal time-lapse imaging of THP1 ASC-GFP (red) stained by LysoView633 (green) and infected by Mtb::BFP (blue).

**Movie S14.** Confocal time-lapse imaging of THP1 mNeonGreen-ALG-2 (green)/Gal3-mScarlet (magenta) infected by Mtb::BFP (blue).

**Movie S15.** Confocal time-lapse imaging of THP1 mNeonGreen-ALG-2 (green)/Gal3-mScarlet (magenta) infected by Mtb::BFP (blue).

**Movie S16.** Simultaneous widefield and TIRF time-lapse imaging of THP1 mNeonGreen-ALG-2 (green)/Gal3-mScarlet (magenta) infected by Mtb::BFP (blue).

**Movie S17.** Widefield time-lapse imaging of THP1 mNeonGreen-ALG-2 (green) stained with Calbryte-590 calcium indicator (grey) and infected by Mtb::BFP (blue).

**Movie S18.** Widefield time-lapse imaging of THP1 Gal3-SNAP(SiR) (magenta) stained with Calbryte-590 calcium indicator (grey) and infected by Mtb::BFP (blue).

**Movie S19.** Confocal time-lapse imaging of THP1 mNeonGreen-ALG-2 (green) in the presence of extracellular propidium iodide (20µg/mL, magenta) and infected by Mtb::BFP (blue).

**Movie S20.** Confocal time-lapse imaging of THP1 mNeonGreen-ALG-2 (green)/Gal3-mScarlet (magenta)/ASC-mIRFP670 (grey) infected by Mtb::BFP (blue).

**Movie S21.** Simultaneous widefield and TIRF time-lapse imaging of THP1 mNeonGreen-ALG-2 (green)/Gal3-mScarlet (magenta) stimulated by silica (100µg/mL).

**Movie S22.** Confocal time-lapse imaging of THP1 mNeonGreen-ALG-2 (green)/Gal3-mScarlet (magenta)/ASC-mIRFP670 (grey) stimulated by silica (100µg/mL).

**Movie S23.** Confocal time-lapse imaging of THP1 mNeonGreen-ALG-2 (green)/Gal3-mScarlet (magenta)/ASC-mIRFP670 (grey) stimulated by LPS (10ng/mL) + nigericin (10µM).

**Movie S24.** Confocal time-lapse imaging of THP1 mNeonGreen-ALG-2 (green)/Gal3-mScarlet (magenta)/ASC-mIRFP670 (grey) stimulated by LPS (10ng/mL) + imiquimod (20µg/mL).

**Movie S25.** Simultaneous widefield and TIRF time-lapse imaging of THP1 mNeonGreen-ALG-2 (green, TIRF and WF)/Gal3-mScarlet (magenta, TIRF and WF)/ASC-mIRFP670 (grey, WF), loaded with TPCS_2a_ (0.4 µg/mL, 16h pulse, 4h chase) and exposed to 405 nm blue light prior to imaging.

## REAGENTS TABLE

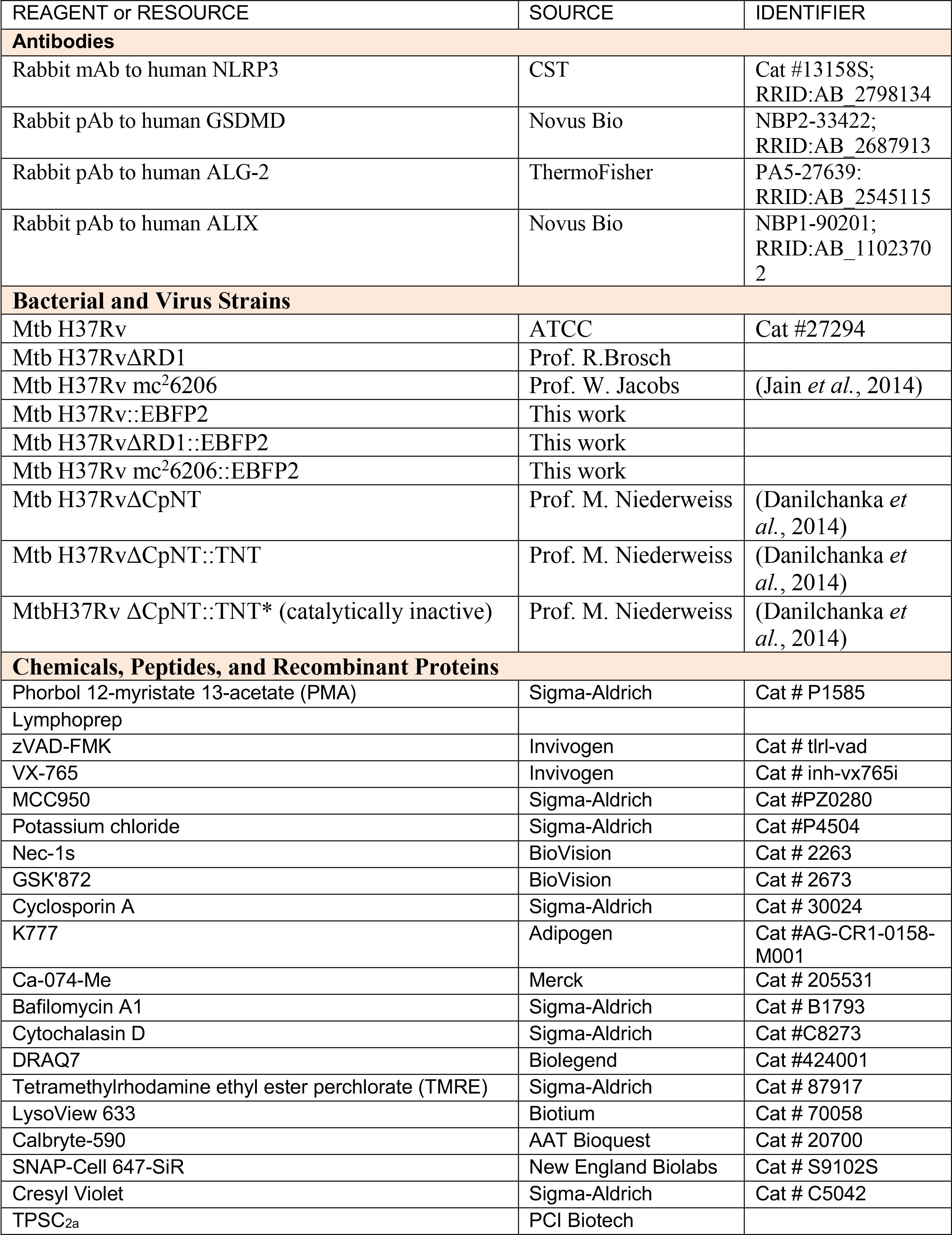
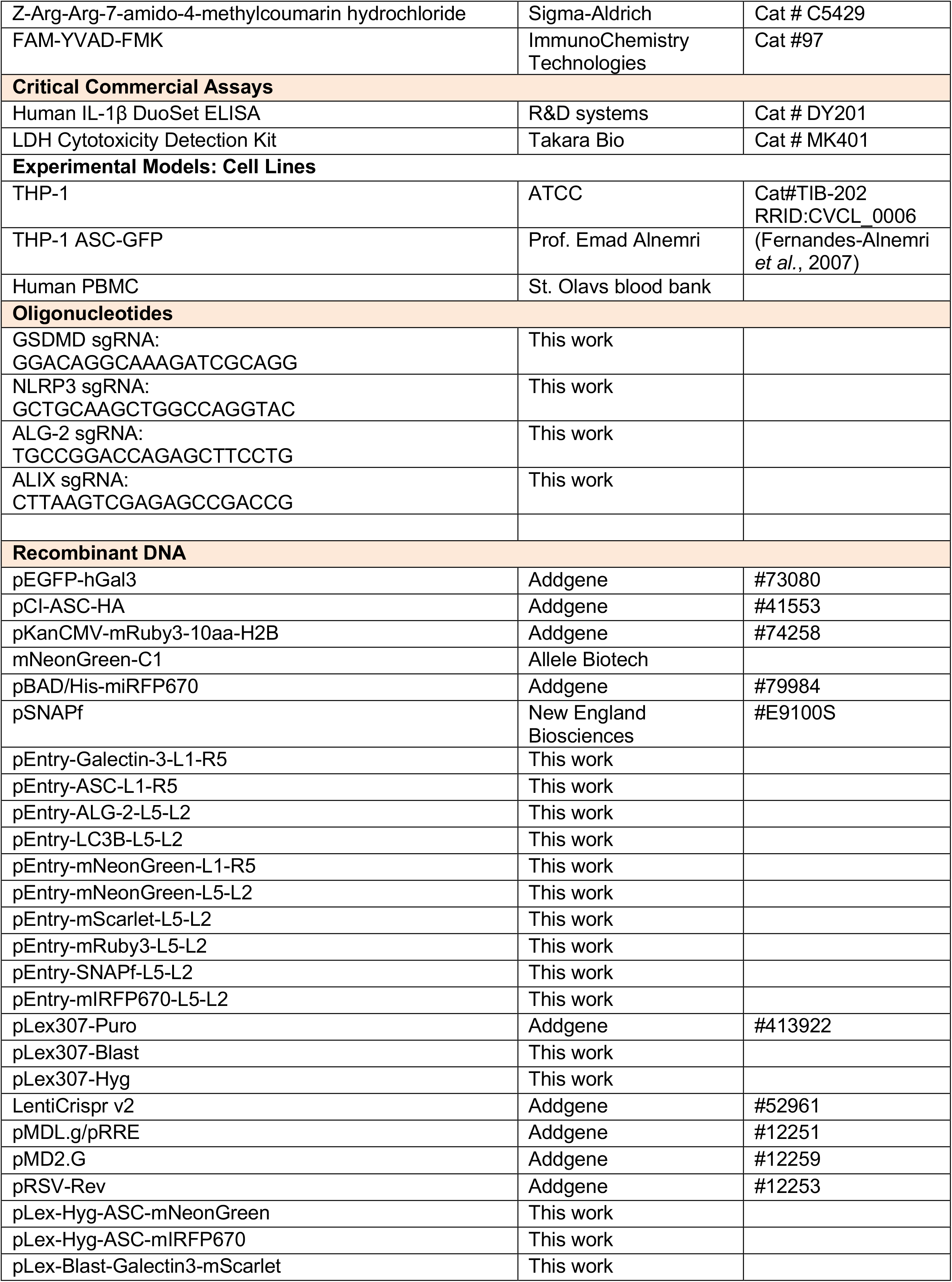
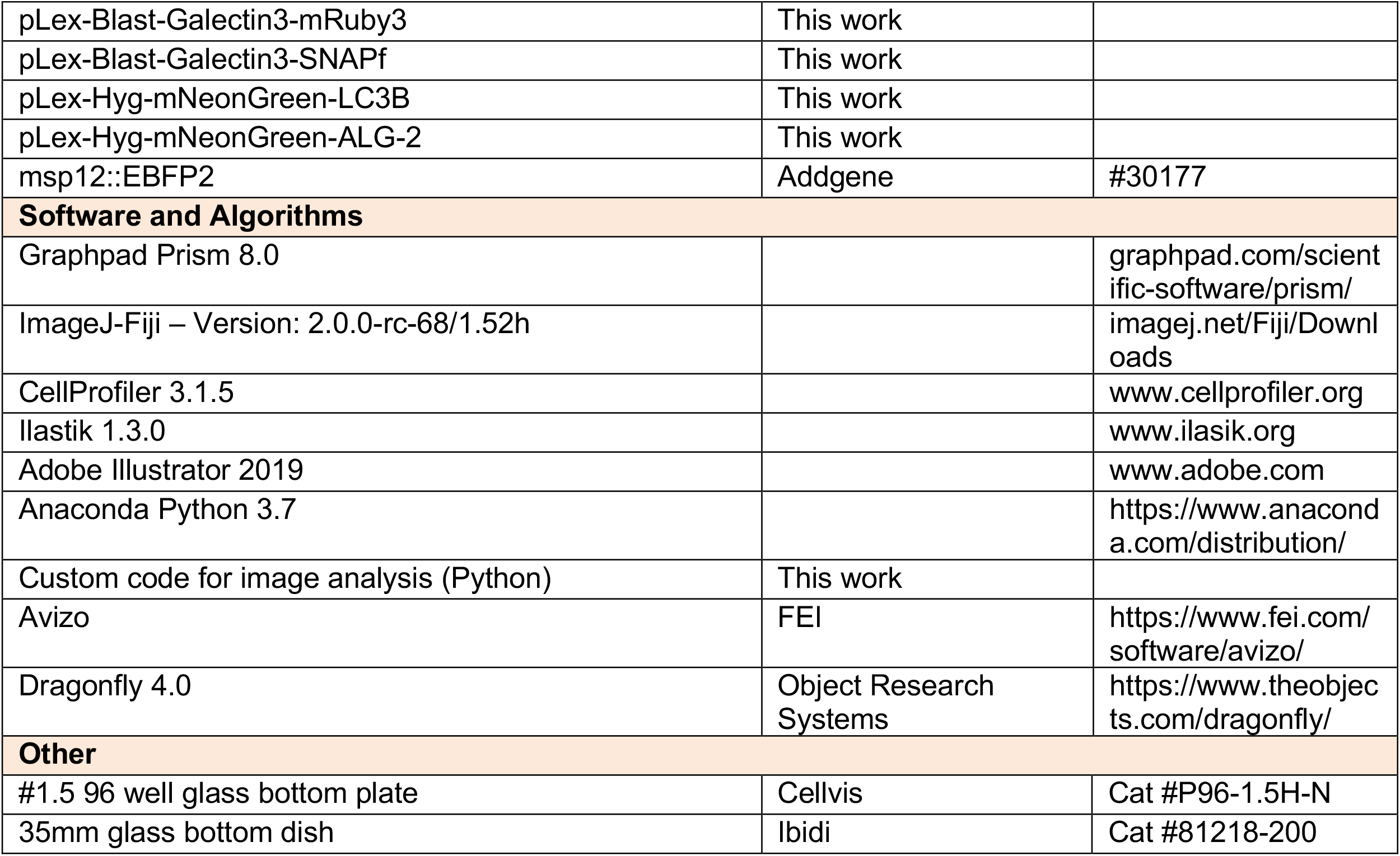

---

**Figure S1, related to main Figure 1.**
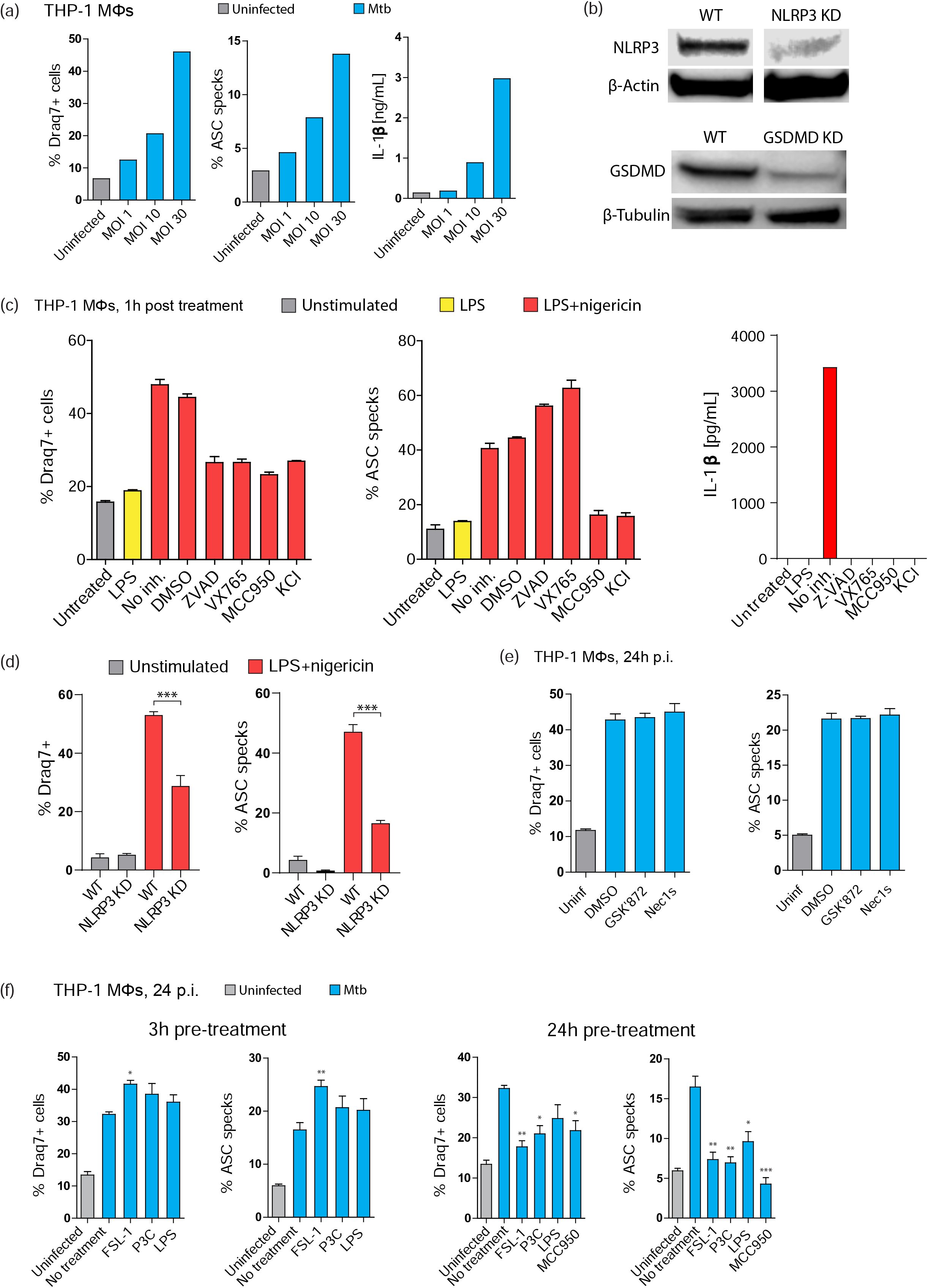
(**a**) THP1 ASC-GFP cells were infected with different MOI of MtbH37Rv. Draq7+ cells, ASC specks and IL-1β release was measured after 24h. Data representative of 2-3 independent experiments. (**b**) THP1 ASC-mNeonGreen cells and THP1 ASC-mNeonGreen depleted of NLRP3 or GSDMD by CRISPR-Cas9 were lysed and protein levels measured by western blotting. (**c**) THP1 ASC-GFP cells were stimulated by LPS (10ng/mL, 3h), not treated or treated with DMSO, Z-VAD-FMK (50µM), VX765 (50µM), MCC950 (10µM) or KCl (40mM) and nigericin (10µM, 1h). Draq7+ cells and ASC specks were quantified from n>2000 cells per condition in triplicate, and IL-1β in the supernatant was quantified by ELISA. (**d**) THP1 ASC-mNeonGreen cells and THP1 ASC-mNeonGreen depleted of NLRP3 by CRISPR-Cas9 were stimulated by LPS and nigericin as above, and Draq7+ cells and ASC specks were quantified from n>2000 cells in triplicates. (**e**) THP1 ASC-GFP cells were treated by DMSO, GSK’872 (5µM) or Nec1s (10µM) and infected by Mtb H37Rv (MOI 20). Draq7+ cells and ASC specks were quantified 24h p.i. for n>2000 cells per condition in triplicate. (**f**) THP1 ASC-GFP cells were untreated or stimulated with FSL-1, Pam3Cys4K or LPS (all 10ng/mL) for 3h or 24h prior to infection with MtbH37Rv and analysed 24h p.i. for Draq7+ cells and ASC specks. n>2000 cells per condition in triplicate were quantified.

**Figure S2, related to main Figure 2.**
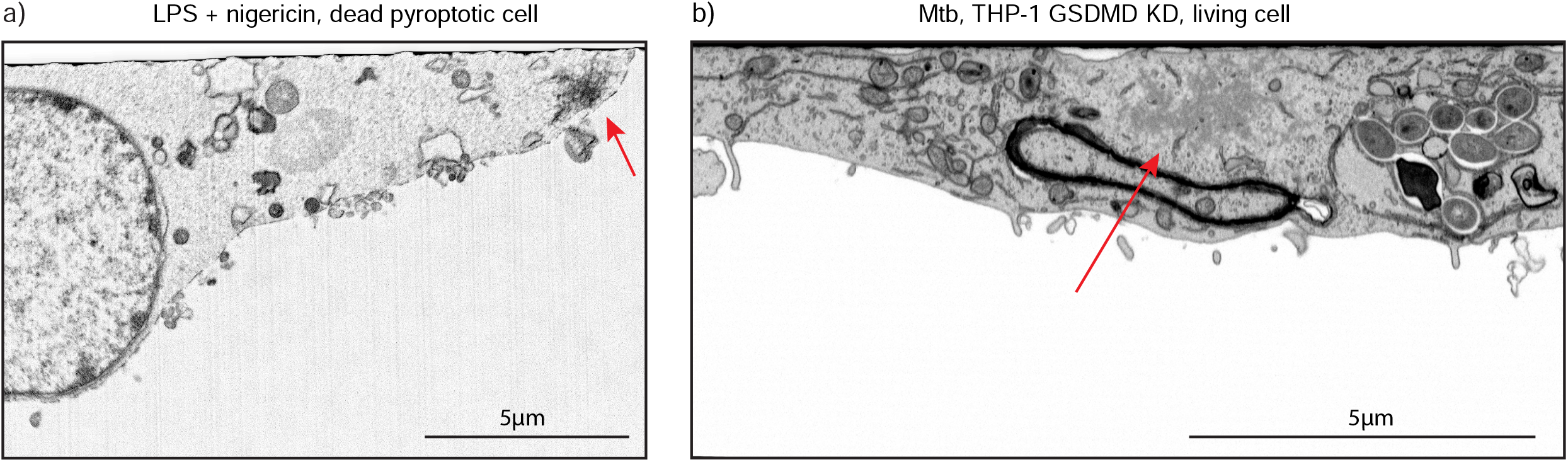
Single slices from FIB-SEM tomography of (**a**) THP1 ASC-GFP cell treated with LPS (10ng/mL, 3h) and nigericin (10µM, 1h) (**b**) THP1 ASC-mNeonGreen depleted of GSDMD by CRISPR-Cas9, infected with Mtb H37Rv and fixed after ASC speck formation while the cell was still alive (DR negative and normal morphology). ASC specks indicated by arrows.

**Figure S3, related to main Figure 3.**
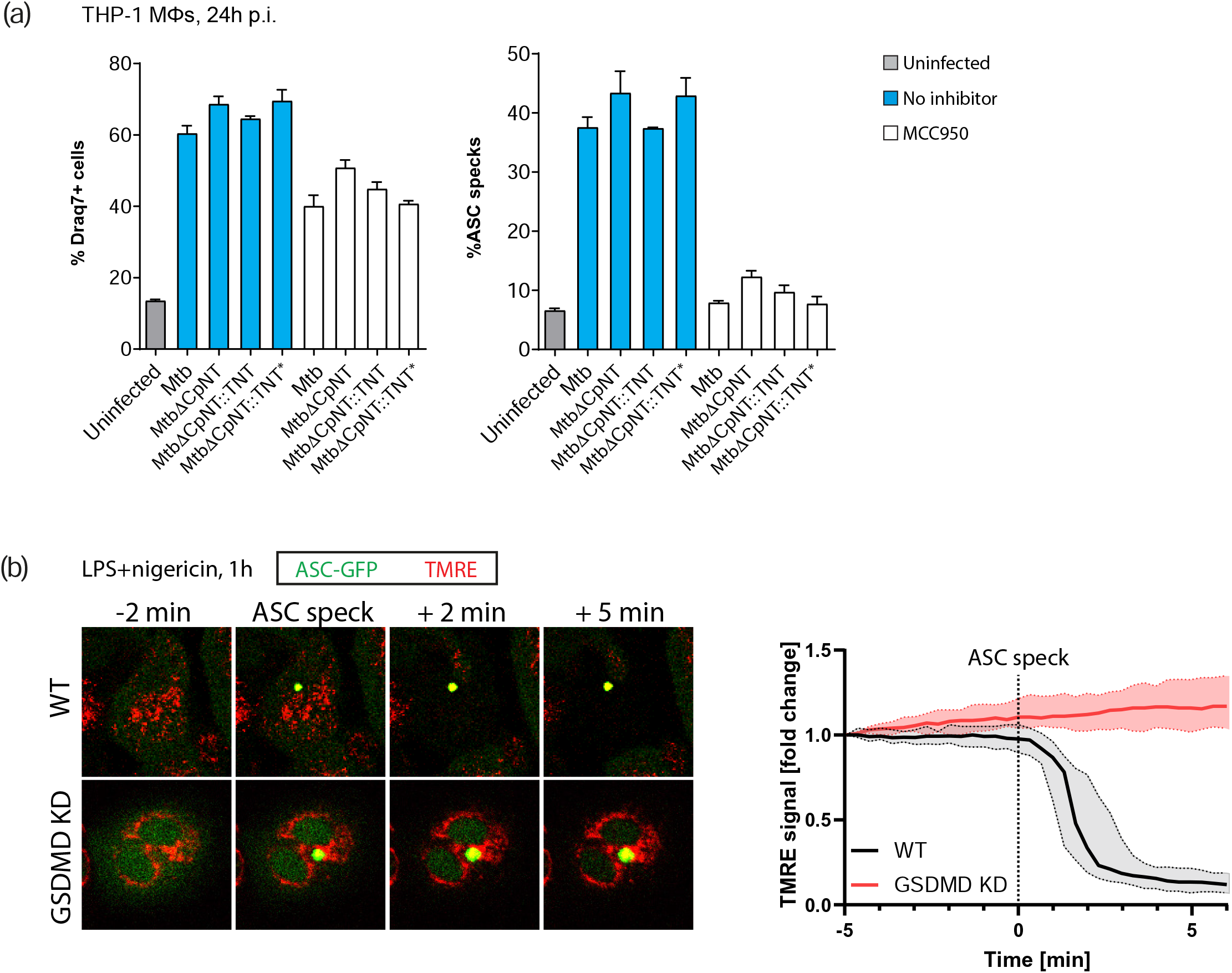
(**a**) THP1 ASC-GFP cells were infected with Mtb H37Rv, Mtb H37RvKCpNT, Mtb H37RvKCpNT::TNT and Mtb H37RvKCpNT::TNT* (catalytically inactive TNT) in the absence or presence of MCC950 (10µM). Draq7+ cells and ASC specks were quantified 24h p.i. for n>2000 cells per condition in triplicate. (**b**) THP1 ASC-GFP cells or THP1 ASC-mNeonGreen cells depleted of GSDMD by CRISPR-Cas9 were labelled with TMRE (10nM) and treated by LPS (10ng/mL) for 3h and nigericin (10µM) and imaged by time-lapse microscopy. Representative images of TMRE and ASC during ASC speck formation are shown. Quantification of TMRE intensity in single cells during ASC speck formation in ASC-mNeonGreen (n=67 cells) and ASC-mNeonGreen GSDMD KD cells (n=44 cells). Median±IQR shown.

**Figure S4, related to main Figure 4.**
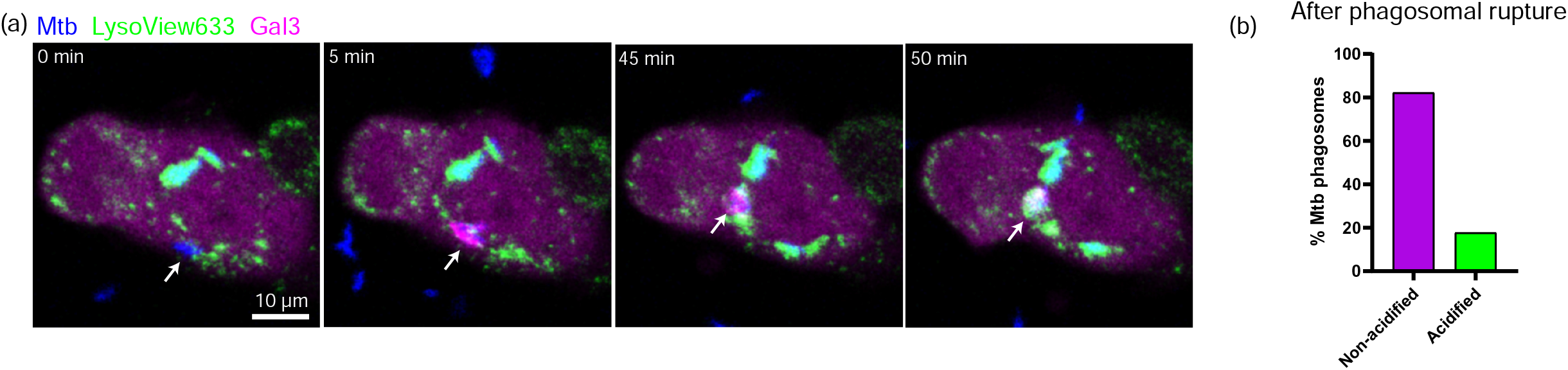
(**a**) Images from confocal time-lapse microscopy of THP1-Gal3-mScarlet cells labelled with LysoView633 and infected by Mtb H37Rv::BFP. (**b**) Proportion of Mtb phagosomes gaining a LysoView signal indicating acidification after a Gal3 event, within 24h or until cell death occurred. n=63 events from one representative experiment.

**Figure S5, related to main Figure 5.**
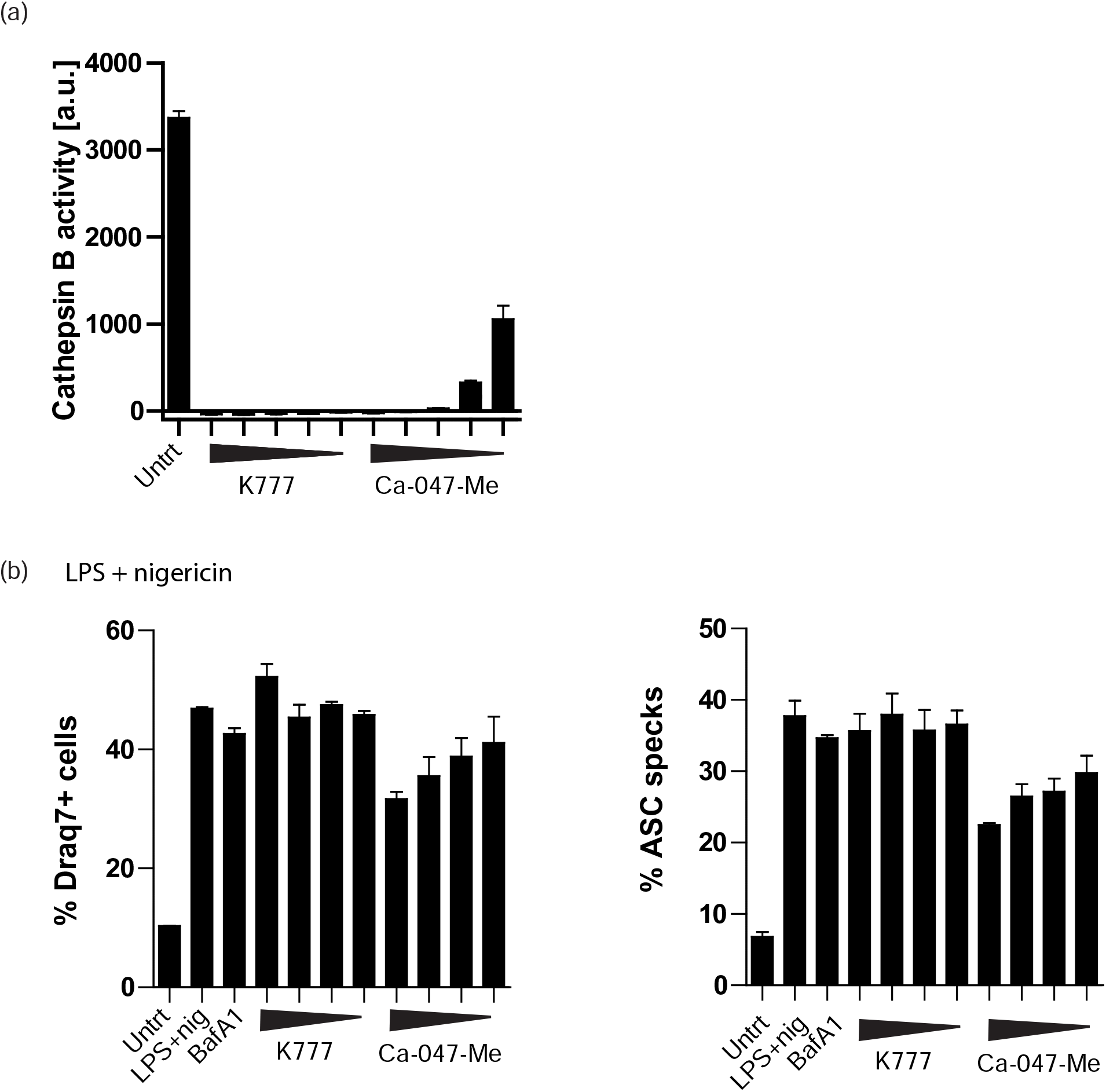
(**a**) Cathepsin B activity measured by Vmax of Z-RR-AMC conversion in total cell lysates without or with 1h pre-treatment by cathepsin inhibitors K777 (30, 15, 7.5, 3.75, 1.6 µM) or Ca-074-Me (30, 15, 7.5, 3.75, 1.6 µM) for 1h. Mean±SEM of technical triplicates. (**b**) Dose response of LPS+nigericin-treated THP-1 macrophages to Bafilomycin A1 (50 nM), K777 (30, 15, 7.5, 3.75 µM) and Ca-074-Me (30, 15, 7.5, 3.75 µM). Cells were primed with 10ng LPS for 3h, pre-treated with inhibitors for 30min than treated with 5µM nigericin for 2h.

**Figure S6, related to main Figure 6.**
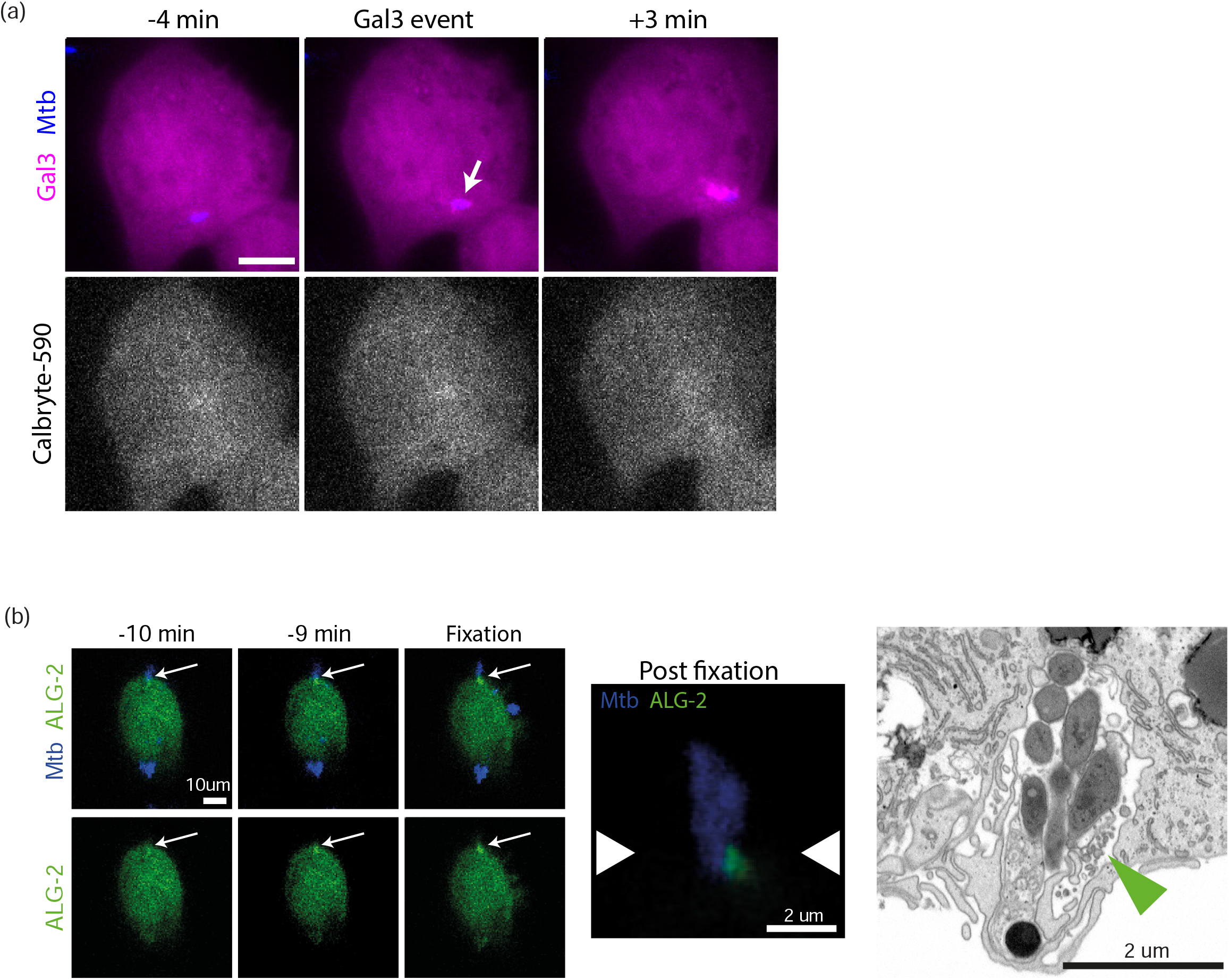
(**a**) Representative images from time-lapse microscopy of Ca2+ sensitive Calbryte-590 signal during an Mtb localized Gal3 event in THP1 Galectin-3-SNAP (SiR) cells. (**b**) Time-lapse confocal imaging of Mtb-localized ALG-2 event indicated by arrow in THP-1 mNeonGreen-ALG-2 cells infected by Mtb H37Rv::BFP. After time-lapse imaging, cells were fixed and imaged by Airyscan and FIB-SEM tomogra-

**Figure S7, related to main Figure 7.**
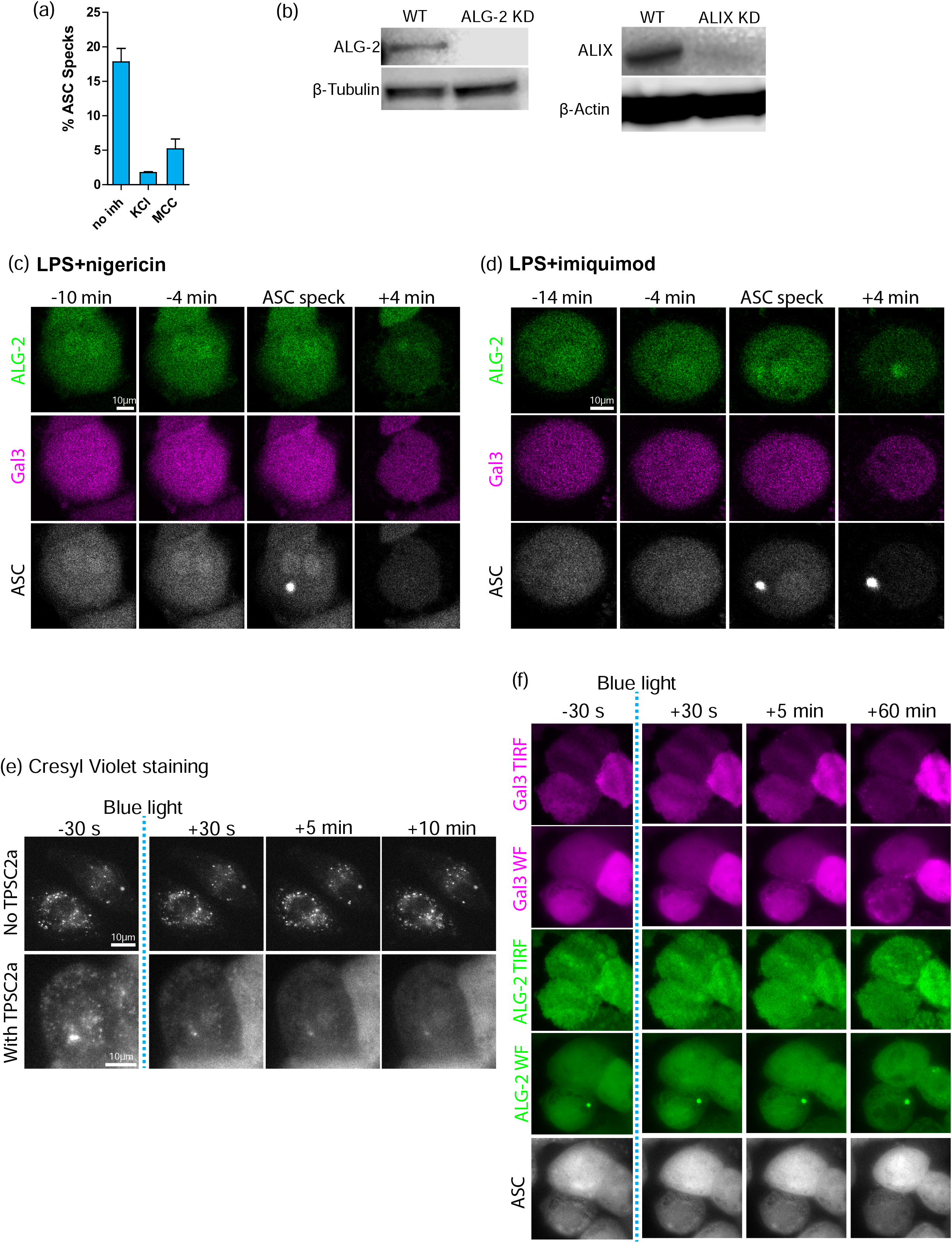
(**a**) Quantification of ASC speck formation from 24h time-lapse experiments of THP1 mNeonGreen-ALG-2/Galectin-3-mScarlet/ASC-mIRFP670 cells not treated or treated by KCl (40mM) or MCC950 (10µM) and infected by Mtb H37Rv::BFP, corresponding to experiment in Figure 7. n>200 cells analysed per condition, mean±SD of 5 fields of view shown. (**b**) ALG-2 or ALIX protein levels in lysates of THP1 ASC-mNeonGreen or ASC-mNeonGreen cells depleted of ALG-2 or ALIX by CRISPR-Cas9. (**c**) THP1 mNeonGreen-ALG-2 (green)/Galectin-3-mScarlet (magenta)/ASC-mIRFP670 (grey) cells treated with LPS (10ng/mL, 3h) and imaged by time-lapse confocal microscopy during treatment with nigericin (10µM). Representative of n=100 similar events. (**d**) THP1 mNeonGreen-ALG-2 (green)/Galectin-3-mScarlet (magenta)/ASC-mIRFP670 (grey) cells treated with LPS (10ng/mL, 3h) and imaged by time-lapse confocal microscopy during treatment with imiquimod (20µg/mL). Representative of n=132 similar events. (**e**) THP1 cells with lysosomes labelled with Cresyl Violet (grey) and pulsed (16h) and chased (4h) with TCSP2a (0.4µg/mL), then exposed to 1s of 10% power 405 laser in a widefield microscope and imaged by time-lapse. Representative of n>50 cells in 5 independent experiments. (**f**) THP1 mNeonGreen-ALG-2 (green)/Gal3-mScalet (magenta)/ASC-mIRFP670 (grey) cells pulsed and chased with TCSP2a (0.4µg/mL), exposed to 405 nm blue light, and imaged for 60 minutes by simultaneous TIRF and widefield time-lapse microscopy. Representative of n>30 cells in 5 independent experiments.

